# Structural serology of polyclonal antibody responses to mRNA-1273 and NVX-CoV2373 COVID-19 vaccines

**DOI:** 10.1101/2024.12.11.628030

**Authors:** Sandhya Bangaru, Abigail M. Jackson, Jeffrey Copps, Monica L. Fernández-Quintero, Jonathan L. Torres, Sara T. Richey, Bartek Nogal, Leigh M. Sewall, Alba Torrents de la Peña, Asma Rehman, Mimi Guebre-Xabier, Bethany Girard, Rituparna Das, Kizzmekia S. Corbett-Helaire, Robert A. Seder, Barney S. Graham, Darin K. Edwards, Nita Patel, Gale Smith, Andrew B. Ward

## Abstract

Current COVID-19 vaccines are largely limited in their ability to induce broad, durable immunity against emerging viral variants. Design and development of improved vaccines utilizing existing platforms requires an in-depth understanding of the antigenic and immunogenic properties of available vaccines. Here we examined the antigenicity of two of the original COVID-19 vaccines, mRNA-1273 and NVX-CoV2373, by electron microscopy-based polyclonal epitope mapping (EMPEM) of serum from immunized non-human primates (NHPs) and clinical trial donors. Both vaccines induce diverse polyclonal antibody (pAb) responses to the N-terminal domain (NTD) in addition to the receptor-binding domain (RBD) of the Spike protein, with the NTD supersite being an immunodominant epitope. High-resolution cryo-EMPEM studies revealed extensive pAb responses to and around the supersite with unique angles of approach and engagement. NTD supersite pAbs were also the most susceptible to variant mutations compared to other specificities, indicating that ongoing Spike ectodomain-based vaccine design strategies should consider immuno-masking this site to prevent induction of these strain-specific responses.

## Introduction

COVID-19 vaccines were developed at an unprecedented pace in the face of a pandemic and were proven efficacious in preventing symptomatic and asymptomatic infections caused by severe acute respiratory syndrome coronavirus 2 (SARS-CoV-2) ^1^. Despite the initial success, the continuous emergence of new SARS-CoV-2 variants, particularly the Omicron lineages, has presented a significant challenge to achieving broad protection against these immune-evasive variants ^2–4^. While administration of either monovalent or bivalent booster shots comprising of newer variants does improve the magnitude and durability of neutralizing antibody responses, achieving durable protective immunity remains a challenge as the virus continues to evolve to evade immunity driven by prior vaccination and/or infection ^5–8^. Design and development of improved vaccines benefit from an in-depth understanding of antigenic and immunogenic properties of the coronavirus (CoV) Spike protein and how it varies with the use of different vaccine platforms. Particularly, structure-guided vaccine design and reverse vaccinology approaches, which have shown promising outcomes in the development of new or improved vaccine candidates against challenging viruses like influenza, HIV, and SARS-CoV-2 ^9–14^, can be applied to preserve the broadly neutralizing epitopes while immune-masking or removing sites that are constantly mutating.

The first generation of FDA-approved (or -authorized for emergency-use) vaccines in the United States utilized various vaccine platforms; Moderna (mRNA-1273) and Pfizer-BioNTech (BNT162b2) are mRNA-based vaccines, Janssen (Ad26.COV2) is a viral vector-based vaccine, while Novavax (NVX-CoV2373) is an adjuvanted recombinant protein vaccine. While they vary in their vaccine constituents, all licensed vaccines in the U.S are based on the two-proline stabilized SARS-CoV-2 Spike protein, the target for protective host antibody responses against the virus ^15–17^. During infection, Spike gets cleaved into two subunits: the S1 subunit that recognizes the human angiotensin-converting enzyme 2 (ACE2) receptor and the S2 subunit that undergoes further cleavage (S2’) and conformational changes to trigger membrane fusion and entry into the host cell ^18–22^. The S1 subunit is comprised of an N-terminal domain (NTD), a receptor-binding domain (RBD) that engages ACE2, and two subdomains (SD1 and SD2) ^20,22^. Isolation of large panels of monoclonal antibodies (mAbs) from vaccinated and naturally infected individuals has revealed that most neutralizing antibodies against the Spike target the receptor-binding domain (RBD) ^17,23–25^. RBD mAbs have been characterized in detail based on their epitopes, gene usage, and mechanisms of neutralization ^26–28^. Neutralizing antibodies to the RBD predominantly bind to or around the receptor-binding motif (RBM) and function by directly or sterically blocking ACE2 receptor engagement and/or inducing premature S1 shedding ^29^. Apart from RBM, RBD mAbs can also bind to core regions of RBD, targeting either the conserved N343 glycan site (outer face) or the cryptic inner face, with a spectrum of binding affinities and neutralizing activity ^30,31^. The other key target for nAbs is the S1 N-terminal domain (NTD) supersite, a glycan-free electropositive surface on the NTD consisting of a mobile β-hairpin and several flexible loops ^25,32–34^. Unlike the RBD, only a few epitopes on the NTD have been associated with neutralizing function, and the exact mechanisms of neutralization remain unclear ^33^. The proposed mechanisms for protection mediated by NTD antibodies include cell-cell fusion inhibition, trimer dissociation, or locking of prefusion Spike conformation as well as Fc-mediated antibody-dependent cellular phagocytosis (ADCP) and antibody-dependent cellular cytotoxicity (ADCC) ^32,33,35^. NAbs towards the more conserved S2 stem helix and fusion peptide regions are also induced by natural infection and vaccination ^36–41^. These mAbs block fusion events and can have remarkable breadth extending across several β-CoVs but are observed at much lower frequencies than S1 antibodies and are typically less potent.

While isolation of mAbs has allowed for detailed characterization of Spike antibodies, our knowledge of the overall epitope landscapes of vaccine-induced polyclonal antibodies (pAb) remains incomplete. Epitope profiling, by either deep mutational scanning or high-resolution linear epitope mapping, has shown that vaccine-elicited antibodies have a broader distribution of epitopes on the RBD compared to those elicited by infection ^42,43^. Another study by Wang et al., compared memory anti-RBD antibodies obtained at 6 months following the initial dose of different vaccine regimens (mRNA, Ad26.COV.2S, ChAdOx1, heterologous ChAdOx/BNT162b2 prime-boost) and discovered significant differences in the RBD epitope distribution between the vaccines tested^44^. These results reveal intrinsic differences in the antigenic properties of SARS-CoV-2 Spike delivered via various vaccine platforms and from natural infections. While these studies have provided key epitope information for vaccine-induced RBD antibodies, there is relatively less knowledge about the other sites on the Spike targeted by vaccines. Moreover, antibody responses to mRNA vaccines have been much better characterized compared to other approved vaccines.

In this study, we investigated antibody responses to both mRNA-1273 and NVX-CoV2373 vaccines using electron microscopy-based polyclonal epitope mapping (EMPEM) analysis, which can provide a comprehensive picture of serum antibody landscapes to the antigen of interest ^45–47^. Both vaccines encode the full-length Spike with two proline substitutions (K986P and V987P) in the S2 region that have been shown to stabilize the Spike in its prefusion conformation and provide better immunogenicity ^20,22,48,49^. Additionally, the Novavax vaccine also contains a 3Q mutation (682-RRAR-685 to 682-QQAQ-685) to confer protease resistance. mRNA-1273 is a lipid nanoparticle (LNP) encapsulated RNA-based vaccine, while NVX-CoV2373 delivers full-length Spike glycoprotein formulated in polysorbate 80 detergent as multivalent nanoparticle immunogens and is adjuvanted with Matrix-M^TM 49,50^. We analyzed serum-derived polyclonal IgGs from both NHPs and clinical trial donors against ancestral Wuhan and variant Spikes to evaluate similarities and differences in the responses between the vaccines and their relative susceptibility to variants. Responses to the immunodominant RBD epitopes in and around the RBM were observed in both NHPs and human donors immunized with either vaccine. Notably, both vaccines elicited a dominant NTD response, mainly targeting either the supersite or the membrane proximal face, with a larger diversity of NTD antibodies seen in the NVX-CoV2373 vaccinated groups. The NTD supersite pAbs were detected in pre-boost and 3 months post-boost samples from clinical trial donors at levels comparable with RBM antibodies and were the most susceptible to variant mutations compared to other specificities.

## Results

### mRNA-1273 and NVX-CoV2373 immunization in NHPs elicit analogous responses

Vaccine responses in NHPs receiving either the NVX-CoV2373 (5 µg) or mRNA-1273 (100 µg) prime-boost series were independently analyzed and reported previously ^49,51,52^. Boosts were administered on days 21 or 28 for NVX-CoV2373 and mRNA-1273, respectively. To investigate the epitope specificities of Spike-reactive pAbs from these NHPs, we performed negative stain EMPEM (ns-EMPEM) with polyclonal Fabs derived from purified serum IgGs from NVX-CoV2373 (Day 35 bleed) or mRNA-1273 (Day 42 bleed) immunized animals. Purified Fabs from four animals from each vaccine group were complexed with a mixture of stabilized SARS-CoV-2 Wuhan (ancestral strain) Spikes and analyzed with ns-EMPEM **(Fig. 1A and B)**. Surprisingly, we observed high levels of Spike trimer dissociation when complexed with pAbs from mRNA-1273 immunized animals while the extent of pAb-induced trimer dissociation was much lower in NVX-CoV2373 immunized samples. Implementing cross-linking of the Spike-pAb complexes following a shorter incubation period with lower pAb-Spike ratios allowed us to recover intact trimer complexes^53^. All animals from both vaccine groups exhibited pAb responses to the RBM (class 1 and class 2), an immunodominant site on the RBD associated with ACE2 blocking activity and potent neutralization. PAbs to the RBD core (classes 3 and 4) were also observed in two NHPs that were immunized with mRNA-1273 but none in the NVX-CoV2373 immunized animals. Class 4 mAbs, like CR3022, that target a cryptic epitope on the RBD core have been previously associated with trimer dissociation properties ^54^ and the presence of such specificities in the mRNA-1273 immunized animals may explain the observed trimer degradation.

**Figure 1.**
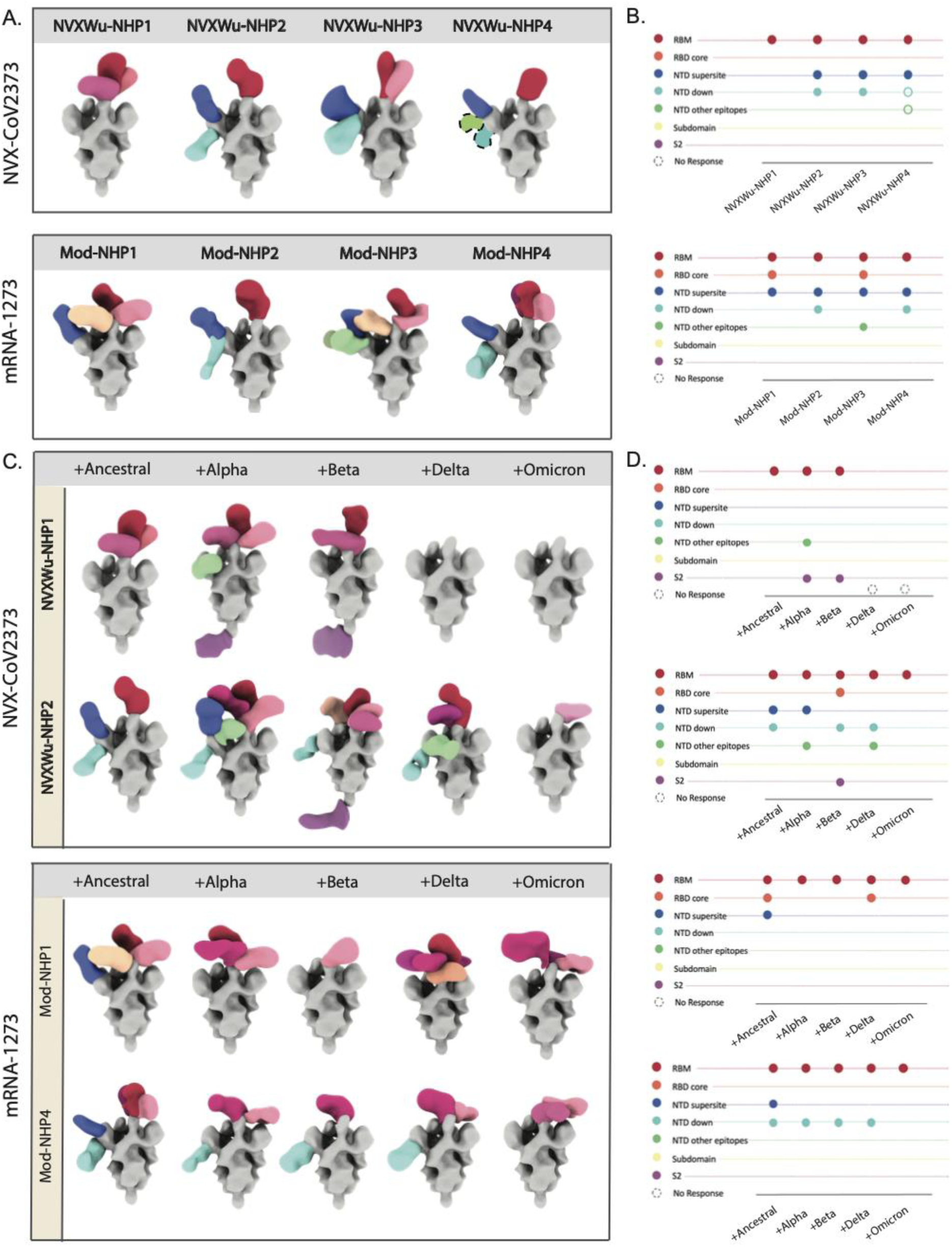
NHP serum pAb responses to NVX-CoV2373 and mRNA-1273 and reactivity to variants. **(A)** Composite maps of Wuhan Spike-Fab complexes and **(B)** their corresponding epitope plots from ns-EMPEM analysis of serum pAbs from NHPs prime-boosted with either NVX-CoV2373 or mRNA-1273 vaccines. **(C)** and **(D)** represent the composite maps of Spike-Fab complexes and their corresponding epitope plots of pAbs from two NHPs from each vaccine group in (A) complexed with Wuhan (ancestral) or variant Spikes. On composite maps, CoV Spikes are colored grey and the Fabs are colored based on epitope specificities as indicated on the Y-axis of epitope plots, and similar specificities are represented by different shades of the indicated color. The open circles on epitope plots correspond to Fab specificities identified by ns-EMPEM 2D class averages but could not be reconstructed in 3D.

The second most prevalent specificity was the NTD supersite with serum from 7/8 NHPs tested containing pAbs targeting this site **(Fig. 1A and B)**. Based on antibody interactions, NTD has five major loops assigned as follows: N1 (residues 14-26), N2 (residues 67-79), N3 (residues 141-156), N4 (residues 177-186), and N5 (residues 246-260), among which the supersite primarily consists of loops N3 and N5 ^32–34^. PAbs to the membrane proximal side of the NTD, hereafter referred to as NTD-down specificity, were also observed in both vaccine groups in 3/4 NVX- CoV2373 and 2/4 mRNA-1273 immunized NHPs. MAbs targeting this region (S2M24 ^33^, DH1052 ^55^, and C1717 ^56^) have been shown to possess varying levels of neutralizing, infection-enhancing, and protective functions. PAbs to other sites on NTD were also seen at lower frequencies, while S2 antibodies were not observed in any animals tested. It is to be noted that the HexaPro (HP) mutations used to stabilize the SARS-CoV-2 Spikes that we employed in our study may preclude binding to the fusion peptide region on S2.

To further investigate if these responses to NVX-CoV2373 and mRNA-1273 based on the Wuhan Spike are cross-reactive to other variants, we tested the pAbs from two NHPs from each immunization group against Alpha (B.1.1.7), Beta (B.1.351), Delta (B.1.617.2), and Omicron (B.1.1.529) Spikes **(Fig. 1C and D)**. Remarkably, certain antibody specificities to the RBM seen in both vaccine groups were able to retain binding against most tested variants, whereas complete abrogation of pAbs binding to the NTD supersite was observed across all variants in both groups. In comparison, NTD-down specificities were less susceptible to variant mutations. Notably, we also observed pAbs to other NTD epitopes and/or to S2 on Alpha, Beta and Delta variants but not against the original Wuhan Spike, indicating that these Wuhan Spike-reactive specificities are indeed present, but their low frequencies or low affinities preclude them from reliable detection by EMPEM **(Fig. 1C and D)**. Factors that can contribute to differences in detection across variants include the overall stability of each variant Spike and the presence of other dominant specificities to the Spike that would affect downstream classification of the low frequency pAbs. In summary, our results reveal RBM and NTD supersite as the most immunodominant epitopes in both vaccine groups with the NTD supersite being highly susceptible to all variant mutations.

### NHP responses to homologous or heterologous prime-boosts with NVX-CoV2373 and Novavax beta Spike immunogen (rS-Beta)

While much of the responses induced by the original prime-boost series target the highly variable regions on the NTD and the RBD, we sought to understand if these responses could be shifted to more conserved epitopes if boosted with a variant Spike instead of the original Wuhan-based immunogen. The Beta variant (B.1.351) was one of the earliest variants of concern (VOC) that possessed several mutations near the NTD supersite (L18F, L242H, 242/244Δ, and R246I) and the RBM (K417N, E484K, and N501Y) that allowed significant immune escape from vaccine-induced responses ^57,58^. To test this, we utilized serum from a Novavax NHP study that compared responses to the original NVX-CoV2373 vaccine, and a beta Spike-based immunogen (rS-Beta) made with a similar formulation ^59^. The study consisted of 3 NHP groups: group 1 receiving a homologous prime-boost of rS-Beta; group 2 receiving a homologous prime-boost of NVX-CoV2373; and group 3 receiving a heterologous prime-boost with NVX-CoV2373 and rS-Beta, respectively. Plasma-derived pAbs isolated from animals within each study group were pooled and complexed with either the Wuhan, Beta, or Delta Spikes. As seen before, ns-EMPEM on the pooled sera from NHPs that received the traditional prime-boost with NVX-CoV2373 contained pAbs targeting the RBM, NTD-supersite, NTD-down, and other NTD epitopes on the Wuhan Spike **(Fig. 2A and B)**. Interestingly, both homologous and heterologous prime-boosts with rS-Beta also exhibited robust responses to Wuhan Spike with a detectable increase in the number of pAb specificities to RBD in NHPs that received the heterologous prime-boost regimen. Additionally, better retention of RBD specificities to Beta and Delta Spikes was apparent in the rS-Beta immunized groups, whereas the loss of NTD supersite responses to variant Spikes were seen across all groups.

**Figure 2.**
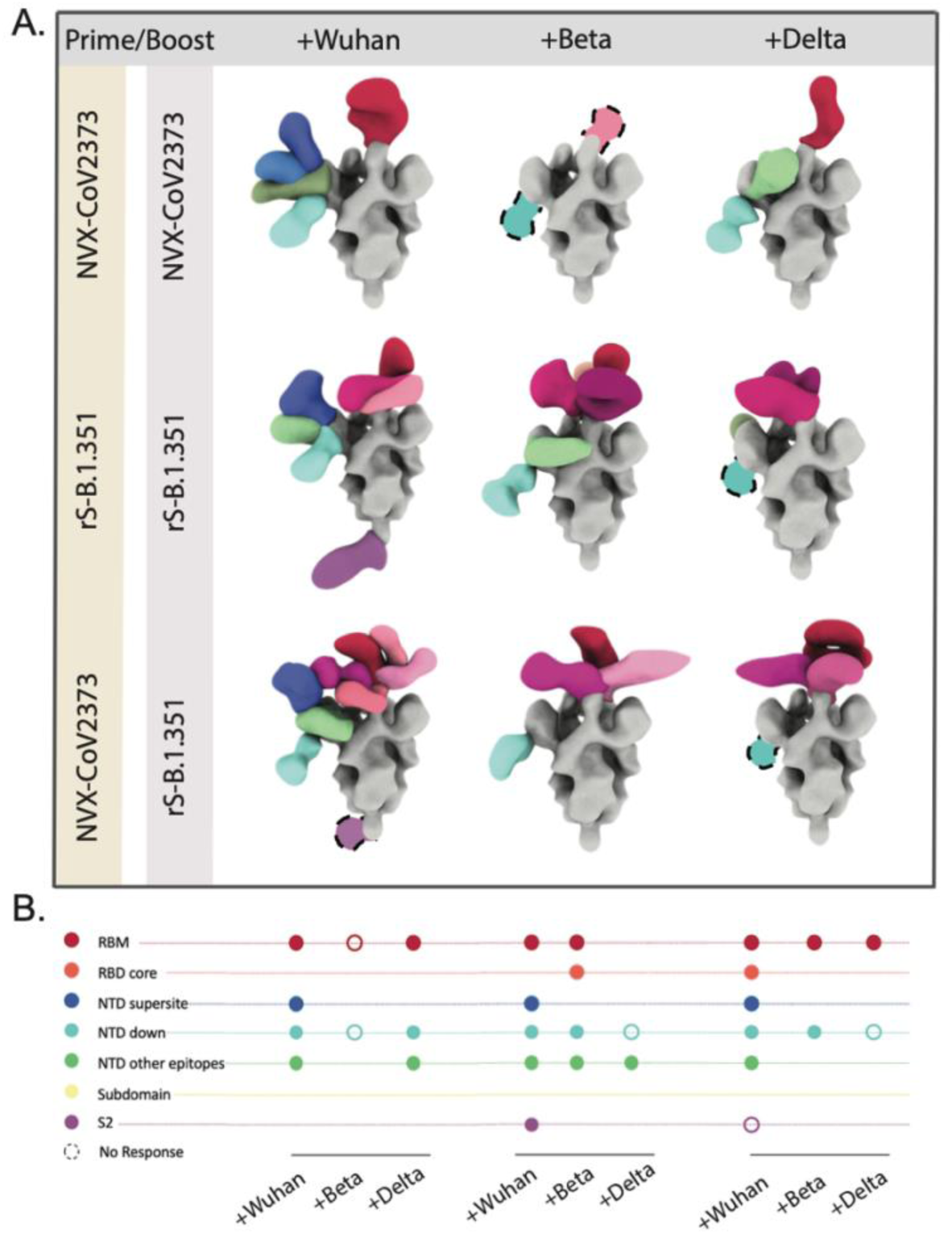
NHP serum pAb responses to homologous or heterologous prime-boost immunizations with NVX-CoV2373 and rS-Beta. **(A)** and **(B)** represent the composite maps of Spike-Fab complexes and their corresponding epitope plots from ns-EMPEM analysis of serum pAbs pooled from four NHPs from each vaccine group complexed with either Wuhan or variant Spikes. CoV Spikes are colored grey and the Fabs are colored based on epitope specificities as indicated on the Y-axis of epitope plots, and similar specificities are represented by different shades of the indicated color. The open circles on epitope plots correspond to Fab specificities identified by ns-EMPEM 2D class averages but could not be reconstructed in 3D.

To examine the molecular basis of pAb responses, we performed cryo-EMPEM on Wuhan Spike complexed with pooled pAbs from NVX-CoV2373 immunized NHPs. Seven Fab-Spike complexes, termed NVX-NHPWu1- NVX-NHPWu7, were resolved at resolutions ranging from 3.8 Å to 4.4 Å, revealing seven distinct epitopes on the NTD **(Fig. 3A, S3, S5 and Tables S1, S2)**.

**Figure 3.**
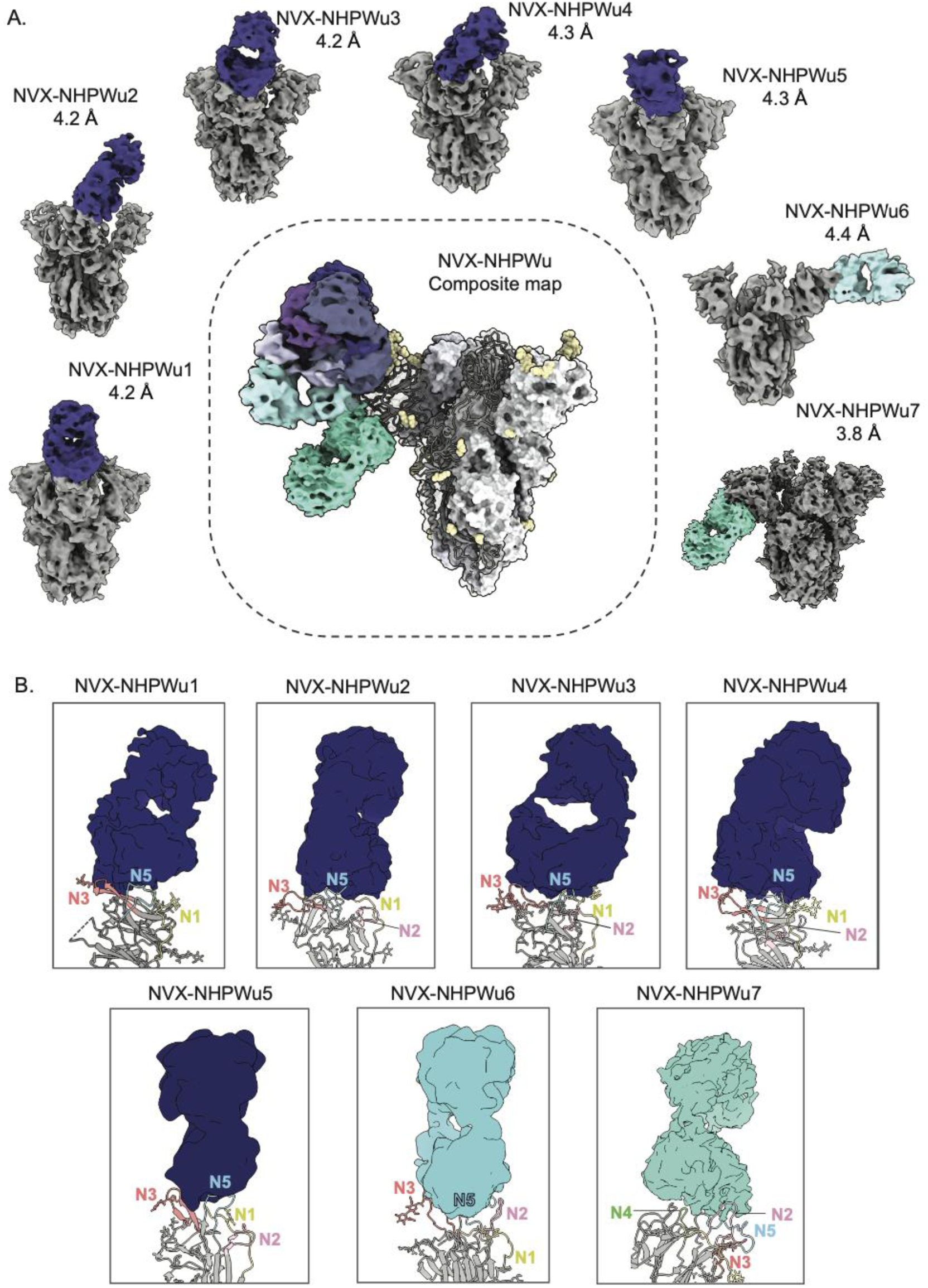
Cryo-EMPEM analysis of polyclonal Fabs from NHPs prime-boosted with NVX-CoV2373. **(A)** Seven cryo-EMPEM reconstructions of Wuhan Spike complexed with distinct polyclonal Fabs derived from pooled sera from NHPs immunized with NVX-CoV2373, prime and boost. The composite map representing all the identified Fab specificities is shown in the center. The supersite antibodies are colored in dark blue, lateral antibodies in light blue and NTD-down antibodies in turquoise. **(B)** Zoomed-in views of Fab-NTD (ribbon representation of docked atomic model) complexes from (A) demonstrating the specific NTD loops being engaged by various pAb specificities. The N1, N2, N3, N4, and N5 loops are colored yellow, pink, orange, green, and blue, respectively. PDBs for docked Spike NTD models, selected based on the closest loop conformations for each map, for NVX-NHPWu1-7 are 7l2f, 7c2l, 7mxp, 7jji, 7sjo, 7mxp, and 7mxp, respectively.

While moderate local resolutions at the epitope-paratope interfaces did not enable atomic model building, docking Spike models into the density allowed analysis of epitopes in detail. Antibodies NVX-NHPWu1-5 all targeted the NTD supersite, interacting largely with the N3 and N5 loops and making additional contacts with either N1 and/or N2 loops **(Fig. 3B)**. The supersite targeted by these antibodies is a large glycan-free surface on the NTD bordered by four glycans at positions N17, N122, N149, and N165, with the N122 glycan being a part of the NVX-NHPWu1 epitope. While these antibodies displayed various angles of approach, they were generally directed downward, facing the viral membrane **(Fig. 3A and 3B)**. Most of these specificities closely resemble published mAbs; NVX-NHPWu1 (mAb Beta-43)^60^, NVX-NHPWu2 (mAb 2–17)^32^, NVX-NHPWu3 (mAb 4–18)^32^, NVX-NHPWu4 (mAb N9)^61^, and NVX-NHPWu5 (mAb A7V3)^62^.

NVX-NHPWu5 belongs to the antibody class (4A8-like antibodies) that is generally encoded by the public clonotype heavy chain VH1-24 gene **(Fig. S4)** ^32,34^. These antibodies typically bind to the supersite loops (N3 and N5 exclusively) primarily with their heavy chain and have a distinct binding angle. NVX-NHPWu6 targets the NTD in a manner distinct from other supersite pAbs, by engaging supersite loops (N3 and N5) but also exhibiting stable interactions with the N2 loop, favored by its lateral angle of approach **(Fig. 3A and 3B)**. The last mAb, NVX-NHPWu7, targets a unique epitope on the membrane proximal face of the NTD, interacting with loops N2, N4, and N5 **(Fig. 3A and 3B)**. The pAb is within proximity to Spike residues 71–74, 184–187, 210–215, and 255–262, and closely resembles the published mAb 2490, an NTD antibody that was shown to enhance infection by inducing an open RBD state required for receptor engagement ^63^.

Next, we examined the pAbs from a single representative NHP that received heterologous NVX-CoV2373/rS-Beta immunization (pAbs complexed with Wuhan Spike) to compare with the original NVX-CoV2373 prime-boost. Samples were not pooled for this study as cryo-EMPEM using pooled NHP pAbs presented challenges for downstream cryo-EMPEM processing (classification of individual Fab-Spike complexes) due to the heterogeneity present in the dataset. A total of eight Fab-Spike complexes (Fabs termed NVX-NHPSA1–NVX-NHPSA8) were resolved at resolutions ranging from 4.0 Å to 4.5 Å **(Fig. 4A, S5, S6 and Tables S1, S3)**. In concordance with NVX-CoV2373 prime-boost responses, the majority of the antibodies targeted the NTD-supersite **(Fig. 4A and 4B)**. Among the supersite antibodies, NVX-NHPSA1, 2, 3, 4, and 7 target an epitope comprising of N3, N5, and N1 but not the N2 loop **(Fig. 4B)**. These antibodies approach the NTD at a downward angle facing the viral membrane while NVX-NHPSA5 and 6 also target the supersite elements N3 and N5 but approach at a lateral angle that allows engagement with the N2 loop (no N1 interactions). Of interest, three of the supersite antibodies (NVX- NHPSA2–4) targeted the beta strands of the N3 beta hairpin rather than the connecting loop, which was less evident in supersite pAbs from homologous prime-boost with NVX-CoV2373 **(Fig. 3B and 4B)**. Notably, most antibodies (NVX-NHPSA2, 3, 5, 6, and 7) induced by heterologous prime-boost did not resemble published mAbs, indicating that boosting with rS-Beta-induced antibodies that engage the NTD in a unique manner. Lastly, NVX-NHPSA8 bound to an epitope similar to that of NVX-NHPWu7 encompassing NTD residues 69–73 (N2 loop), 184–187 (N4 loop), 210– 215, and 257–262 (N5 loop) **(Fig. 3B and 4B)**. Taken together, our results indicate that while both homologous and heterologous prime-boost regimens with NVX-CoV2373/rS-Beta commonly elicit responses to the immunodominant supersite and the membrane proximal face of NTD, boosting with rS-Beta target may exhibit a shift in the epitope specificities towards the beta strands on the N3 hairpin structure.

**Figure 4.**
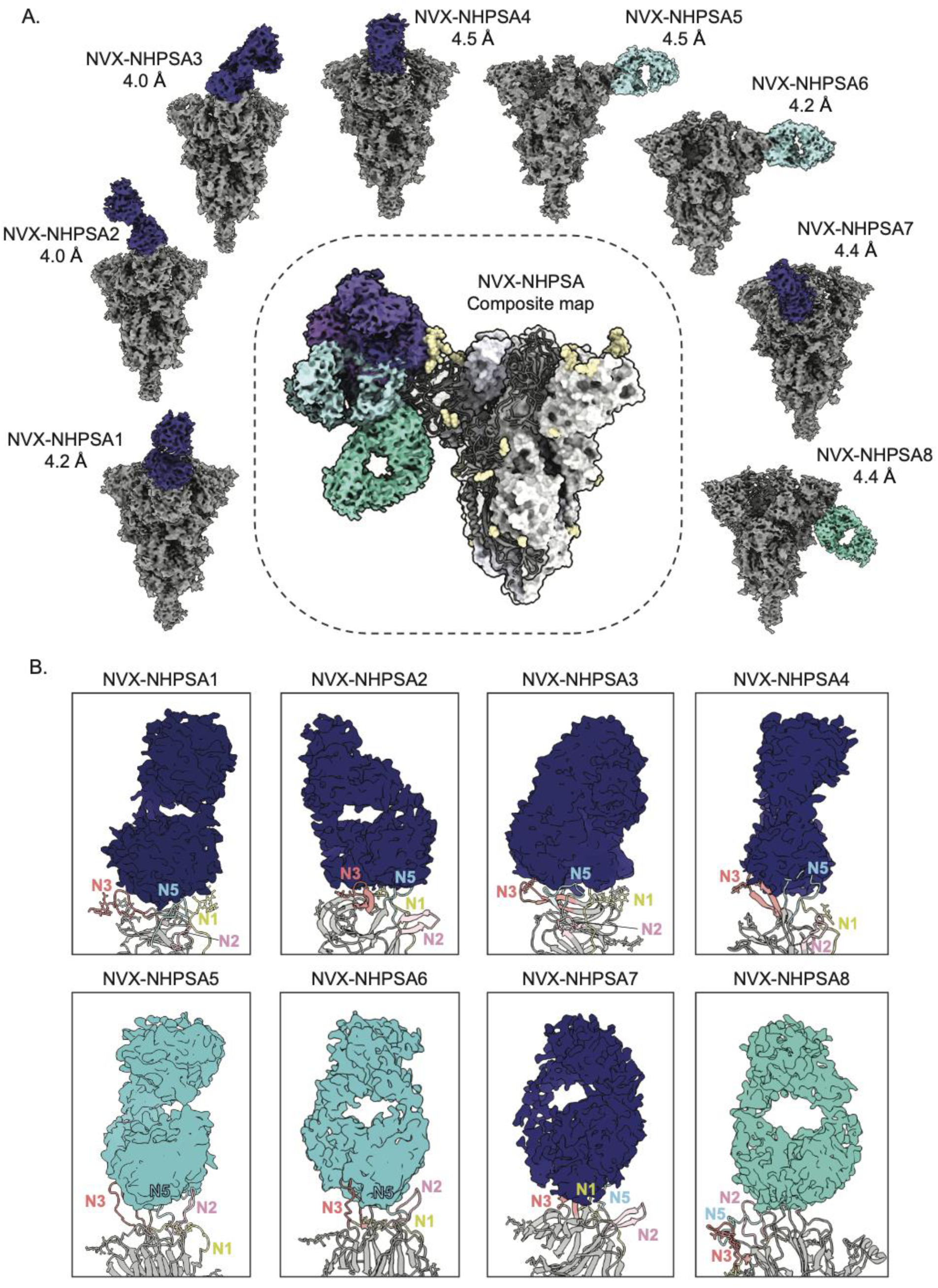
Cryo-EMPEM analysis of polyclonal Fabs from NHP that received heterologous prime-boost immunizations with NVX-CoV2373 and rS-Beta. **(A)** Eight distinct cryo-EMPEM reconstructions of Wuhan Spike complexed with polyclonal Fabs derived from an NHP that was primed with NVX-CoV2373 and boosted with rS-Beta. The composite map representing all the identified Fab specificities is shown in the center. The supersite antibodies are colored in dark blue, lateral antibodies in light blue and NTD-down antibodies in turquoise. **(B)** Zoomed-in views of Fab-NTD (ribbon representation of docked atomic model) complexes from (A) demonstrating the specific NTD loops being engaged by various pAb specificities. The N1, N2, N3, N4, and N5 loops are colored yellow, pink, orange, green, and blue, respectively. PDBs for docked Spike NTD models, selected based on the closest loop conformations for each map, for NVX-NHPSA1-8 are 7mxp, 7l2c, 7jji, 7sj0, 7mxp, 7mxp, 7l2d, and 7mxp, respectively.

### EMPEM analysis of clinical trial donors reveal slight differences in antigenicity across vaccine platforms

Next, we investigated the antibody responses in clinical trial donors who received either the NVX-CoV2373 or mRNA-1273 vaccine series. PAbs were isolated from serum samples collected at day 29 (Mod-D29) and day 43 (Mod-D43) from mRNA-1273 vaccinees, and day 21 (Nvx-D21), day 49 (Nvx-D49), and day 105 (Nvx-D105) from NVX-CoV2373 vaccinees. EMPEM was performed with purified Fabs from four NVX-CoV2373 and five mRNA-1273 donor samples, each complexed with a SARS-CoV-2 Spike mix **(Fig. 5A and 5B)**. Analysis of responses at the pre-boost time points, Mod-D29 and Nvx-D21, showed early elicitation of RBM, NTD supersite, and NTD-down antibodies in mRNA-1273 vaccinees, while the overall responses detected in NVX-CoV2373 vaccinees were limited. 3D reconstructions of Fab-bound Spikes were possible only for 3/5 NVX-CoV2373 donors; NTD supersite and RBM antibodies were detected in three donors and one donor, respectively **(Fig. 5A)**. Notably, the pre-boost bleeds for NVX- CoV2373 were collected at day 21 rather than day 29, which may correspond to the lower levels of antibodies detected for Nvx-D21. Nevertheless, NTD supersite antibodies, along with RBM antibodies, were the major specificities elicited post-prime in both vaccine groups.

**Figure 5.**
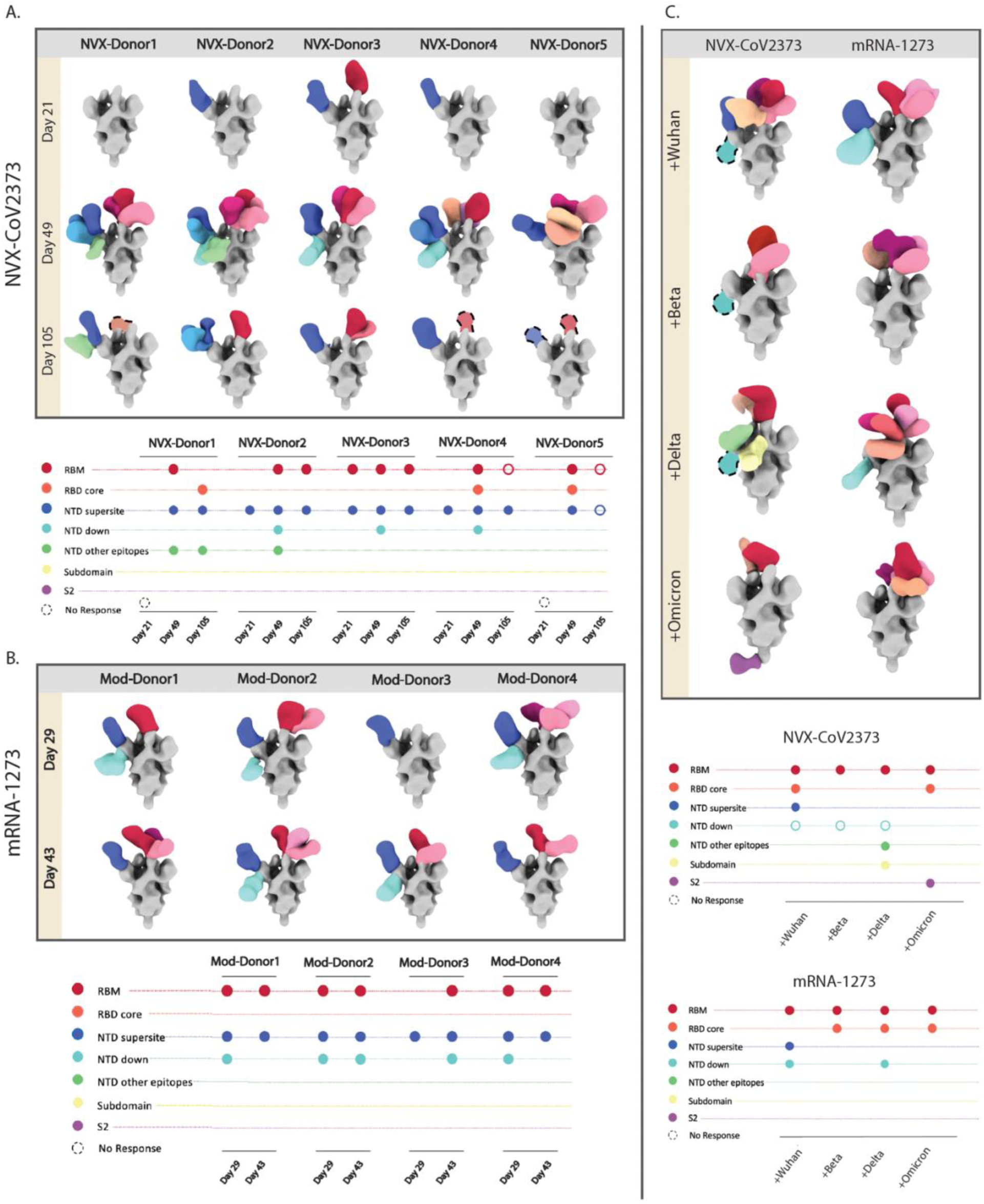
EMPEM analysis of pAbs in clinical trial donors that received either NVX-CoV2373 or mRNA-1273 vaccine series. Composite maps of Wuhan Spike-Fab complexes and their corresponding epitope plots from ns-EMPEM analysis of pAbs derived from sera collected at **(A)** three time points (Days 21, 49, and 105) from five donors who received NVX-CoV2373 vaccine and at **(B)** two time points (Days 29 and 43) from four donors who received mRNA-1273 vaccine. **(C)** Ns-EMPEM analysis of pooled serum pAbs from donors prime-boosted with either NVX-CoV2373 (Day 49) or mRNA-1273 (Day 43) in complex with Wuhan- or variant-derived Spikes. On composite maps, CoV Spikes are colored grey and the Fabs are colored based on epitope specificities as indicated on the Y-axis of epitope plots, and similar specificities are represented by different shades of the indicated color. The open circles on epitope plots correspond to Fab specificities identified by ns-EMPEM 2D class averages but could not be reconstructed in 3D.

The NVX-CoV2373 post-boost samples, Nvx-D49, represented a more diverse epitope profile targeting additional sites on NTD and the RBD core in addition to RBM and NTD supersite while Mod-D43 had similar specificities as Mod-D29 pre-boost **(Fig. 5A and 5B)**. NTD-down responses were observed in half of the donors (Nvx-D49 and Mod-D43) followed by RBD core antibodies, which were only detected in NVX-CoV2373 donors. Importantly, NVX-CoV2373 vaccinees had a larger diversity of NTD pAb specificities compared to mRNA-1273 vaccinees, including pAbs to the supersite and pAbs targeting NTD at lateral angles, potentially engaging the N2 loop, as evidenced by cryo-EMPEM analysis of NVX-CoV2373 immunized NHPs **(Fig. 5A, 5B, and 3B)**. The responses at the Nvx-D105 (∼3 months post-boost) time point were considerably lower, in line with reports of vaccine-induced antibodies undergoing rapid decay until they reach a steady state ^64,65^. Although there was an overall reduction in antibodies to all epitopes, the NTD supersite specificities were retained in all donors along with certain RBM responses, while the NTD-down and RBD core responses were not detected in any donor at this time point.

Next, we investigated the variant reactivity of antibodies at Mod-D43 and Nvx-D49 samples by pooling polyclonal Fabs from multiple donors at these time points and performing ns- EMPEM with Spikes from either the ancestral Wuhan or variant Beta, Delta, or Omicron strains **(Fig. 5C)**. As seen with NHP variant analysis, there was a complete abrogation of NTD supersite antibodies across all variants in both vaccine groups. Loss of NTD-down specificity was also observed against the Omicron variant Spike for both groups. Notably, several RBD antibodies retained binding to all the tested variants in both groups, with the Mod-D43 pool exhibiting a higher diversity of RBD responses to each variant Spike compared to the Nvx-D49 pool. Taken together, findings from clinical trial donor sera highlight some differences in antigenicity between NVX-CoV2373 and mRNA-1273 and the overall susceptibility of NTD supersite antibodies to variant mutations.

### CryoEMPEM of mRNA-1273 vaccinated donors reveal diverse responses to the NTD supersite

To further investigate the molecular details of antibody epitopes elicited by the mRNA-1273 vaccine, cryo-EMPEM was performed on Wuhan Spike complexed with pooled Mod-D43 pAbs. Eight high-resolution maps (3.2 Å to 3.8 Å) of Spike-Fab complexes were reconstructed from this dataset; two recognizing the RBM, five targeting the NTD-supersite and one to the NTD membrane proximal face **(Fig. 6A, S8, S11 and Tables S1, S4)**. Both RBM antibodies (ModWu- RBD1 and ModWu-RBD2) targeted the class 1 site on RBD and displayed nearly identical epitope-paratope features, spare the differences in the RBD conformations between the reconstructions **(Fig. 6A and 6B)**. Comparison of published RBD mAbs with ModWu-RBD1 density revealed a close resemblance with mAb P4A1 (PDB 7CJF), a potent antibody that functions by directly blocking ACE2 receptor engagement ^66^. Docking of the P4A1-RBD structure into the ModWu-RBD1 density showed similarities in the epitope-paratope interface, including overlap with the ACE2 binding site as well as highly overlapping HCDR lengths and conformations **(Fig. 6B)**.

**Figure 6.**
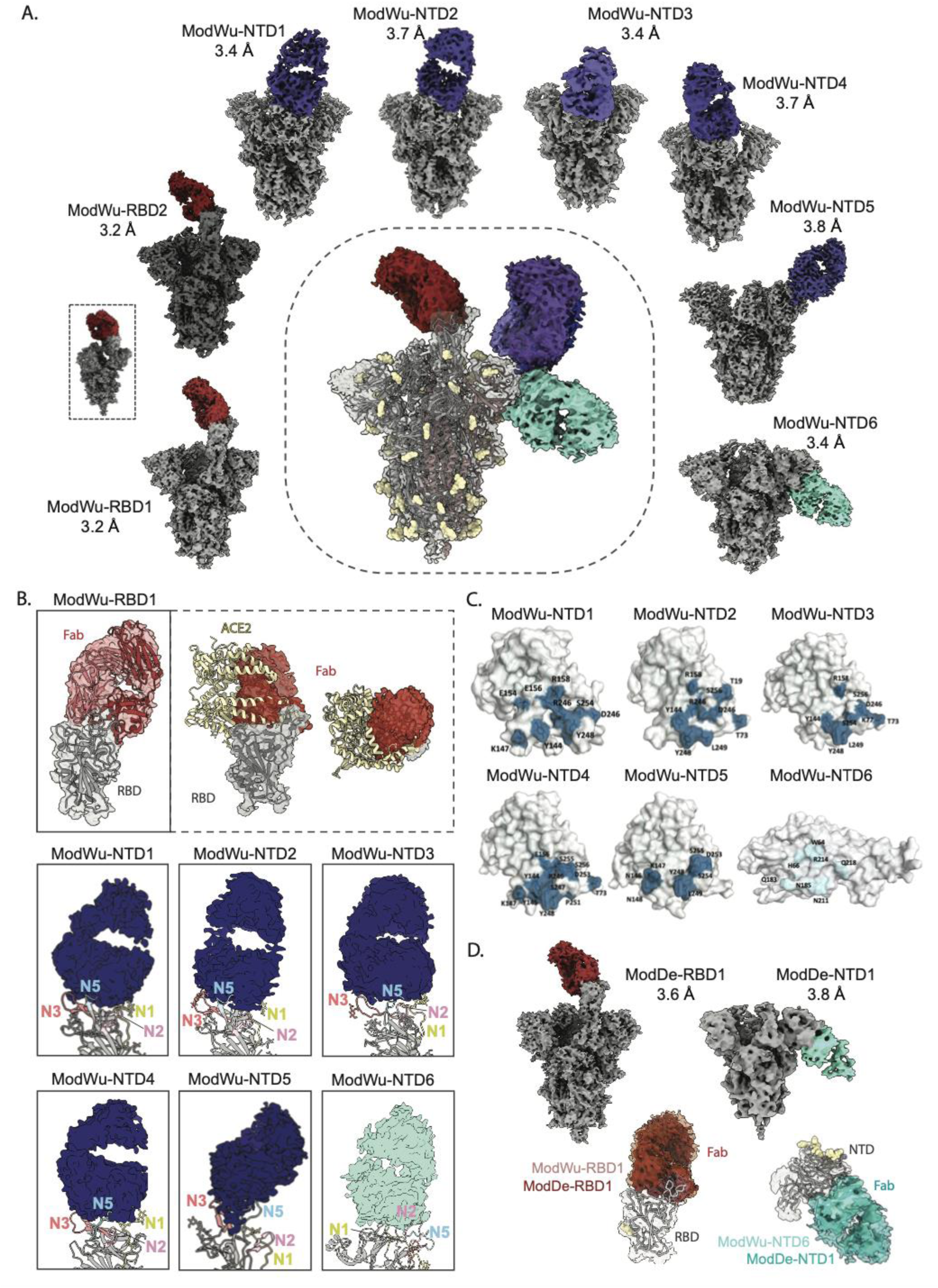
Cryo-EMPEM analysis of polyclonal Fabs from donors vaccinated with mRNA-1273. **(A)** Eight cryo-EMPEM reconstructions of Wuhan Spike complexed with distinct polyclonal Fabs, two RBD and six NTD, derived from pooled sera from mRNA-1273 vaccinees. The composite map representing all the identified Fab specificities is shown in the center. The RBM Fabs are colored in red, NTD supersite Fabs in dark blue and NTD-down Fab in turquoise. **(B)** Zoomed-in views of Fab-RBD and Fab-NTD (ribbon representation of docked atomic model) complexes from (A), demonstrating the epitope specificities of various Fabs. The ModWu-RBD1 map, docked with the atomic model of the P4A1-RBD complex (7cjf), shows binding to RBM class 1 epitope that overlaps with ACE2 RBS. For ModWu-NTD1-6 maps, the NTD loops N1, N2, N3, N4, and N5 are colored yellow, pink, orange, green, and blue, respectively. PDBs for docked NTD models, selected based on the closest loop conformations for each map, for ModWu-NTD1-6 are 7jji, 7jji, 7mxp, 7l2e, 8swh, and 7mxp, respectively. **(C)** Surface representation of the NTD displaying residues in the epitope–paratope interface, identified by MD simulations. **(D)** Two cryo-EMPEM reconstructions of Delta Spike complexed with distinct polyclonal Fabs, one RBD and one NTD, derived from pooled sera from mRNA-1273 vaccinees used in (A). Bottom panel in (**D**) is the overlay of Fab-delta Spike complexes, ModDe-RBD1 and ModDe-NTD1 (solid color), with Fab-Wuhan Spike complexes (transparent surface), ModWu-RBD1 and ModWu-NTD6, respectively.

The five NTD supersite antibodies (ModWu-NTD1-5) targeting the N3 and N5 loops fell into three broad classes based on their epitopes **(Fig. 6B)**. In addition to engaging the supersite elements N3 and N5 loops, pAbs ModWu-NTD1 and ModWu-NTD2 made N74 glycan contacts, while pAbs ModWu-NTD3 and ModWu-NTD4 engaged the N17 glycan on the N1 loop with different angles of approach. ModWu-NTD5 interacted exclusively with the N3 and N5 loops, similar to pAb NVX-NHPWu5, belonging to the 4A8-like antibody class. As seen with responses from NVX-CoV2373 immunized NHPs, pAbs ModWu-NTD2, ModWu-NTD3, ModWu-NTD4, and ModWu-NTD5 resemble published NTD mAbs 2–17^32^, S2M28^33^, 4–18^32^, and TXG-0078^67^, respectively **(Fig. S9)**. The only non-supersite antibody, ModWu-NTD6, bound to the membrane-proximal face of the NTD near the residue ranges 24–30, 68–76, 184–188, 209–216, and 251–263. Due to the high epitope variability of the NTD, we performed molecular dynamics (MD) simulations to capture conformational changes of this supersite upon antibody binding. Within our simulations, we observe multiple conformational states of loops (N1, N3, and N5). By combining the different conformational states with the available high resolution cryo-EMPEM maps, we were able to identify regions/residues within the NTD supersite that are likely to contribute to antibody binding and recognition. ModWu-NTD1-5 targeting the supersite revealed substantial overlaps in the predicted interaction sites, while ModWu-NTD6 recognition of the NTD is mediated by residues distinct from the supersite. Overall, the epitopes targeted by these pAbs were analogous to those identified in the NVX-CoV2373 immunized NHPs. However, we did not detect any specificities targeting the NTD at a lateral angle favoring N2 loop interactions, which were identified in both the NVX-CoV2373 (NVX-NHPWu6) and NVX-CoV2373/rS-Beta (NVX- NHPSA5 and NVX-NHPSA6) immunized animals in addition to the ns-EMPEM analysis of NVX-CoV2373 vaccinees, indicating some antigenic differences between the two vaccine platforms.

Next, we investigated the same Mod-D43 pAb pool in complex with the Delta Spike using cryo-EMPEM to identify the Fab specificities that retain binding to the Delta Spike **(Fig. 6C, S10, S11 and Tables S1, S5)**. Two Fab-Spike (Delta) complexes, ModDe-RBD1 and ModDe-NTD1, were reconstructed to resolutions of 3.6 Å and 3.8 Å, respectively. ModDe-RBD1 pAb interacted with the RBD apex in a manner similar to ModWu-RBD1. Docking the density for ModDe-RBD1 into ModWu-RBD1 revealed an identical angle of approach and engagement, suggesting that these antibodies belong to the same representative pAb family. ModDe-NTD1 targeting the membrane proximal face of the NTD also aligned well with ModWu-NTD6, however the local resolutions in the Fab-NTD region were insufficient to make further conclusions. Remarkably, we were not able to reconstruct any of the supersite antibodies to the Delta Spike, confirming the complete loss of this specificity to variant Spikes and further validating our findings from ns-EMPEM analyses.

Collectively, cryo-EMPEM analyses of sera across all vaccines tested in this study reinforce the overall immunodominance of the NTD supersite, as evidenced by the wide diversity of responses targeting this site. Previous structural analysis of SARS-CoV-2 Spike NTDs, both in apo and antibody-bound states, have shown that NTD supersite elements, N3-beta hairpin, and N1 and N5 loops can exist in multiple conformations to accommodate antibody binding ^32–34,48^. An overview of conformational changes in these loops observed in our MD simulated models, as well as published NTD structures of Spike-antibody complexes, demonstrate the plasticity of these elements contributing to extensive epitope variability **(Fig. 7A and 7B)**. In addition to being a glycan-free surface, the highly flexible and disordered loops encompassing the NTD supersite may provide an important factor for eliciting strong antibody responses, as evidenced in this study.

**Figure 7.**
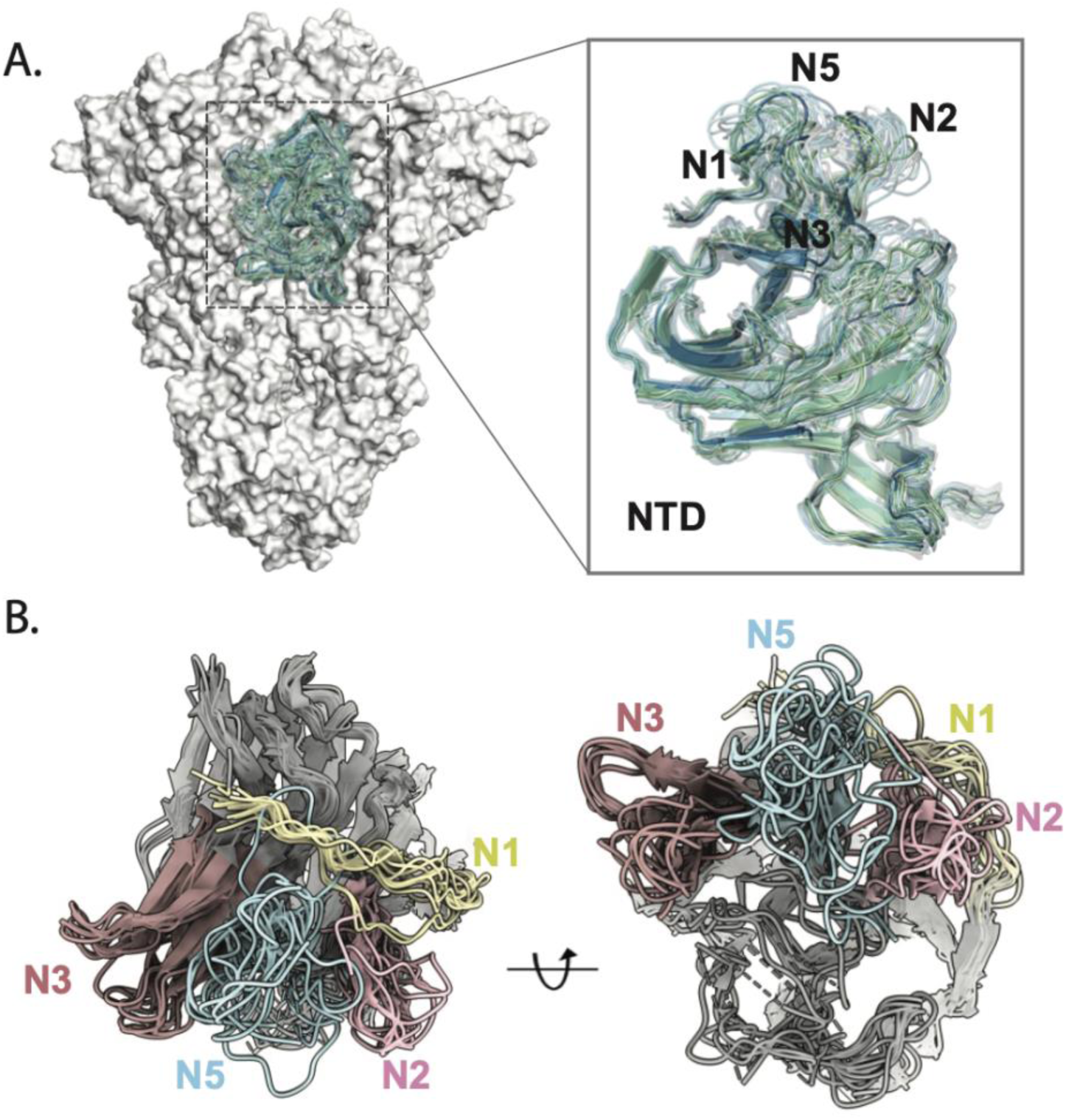
Conformational flexibility of NTD. **(A)** NTD conformations upon antibody binding captured by MD simulations and **(B)** ribbon representation overlay of NTDs from published atomic models displaying the plasticity of loops N1 (residues 14–26), N2 (residues 67–79), N3 (residues 141–156), and N5 (residues 246–260).

## Discussion

Mapping the epitope profiles of vaccine-induced antibodies not only provides valuable information for structure-guided design approaches but can be a critical component for assessing vaccine functionality, predicting vaccine efficacy against emerging variants, and evaluating shifts in antigenicity in the post-vaccination era. Here, we evaluated the antigenic landscapes of Spike proteins delivered using two different vaccine platforms in immunized NHPs and clinical trial donors. Overall, we observed similar responses to both vaccines in NHPs and clinical trial donors, with slight differences in the diversity of responses to RBD and NTD across both vaccines. RBM, class 1 and 2 antibodies have been previously shown to be induced early after infection or mRNA- immunization ^68^ while Ad26.COV.2S vaccination was shown to induce more RBD core, class 3 and 4, early memory B cell responses ^69^. Our EMPEM analysis likewise revealed dominance of RBM pAbs over RBD core antibodies in both mRNA-1273 and NVX-CoV2373 vaccinated donors, pre- and post-boost. Notably, NHPs that received mRNA-1273 had increased prevalence of class 3/4 pAbs than clinical trial donors and NHPs immunized with NVX-CoV2373. Cryo- EMPEM of mRNA donor sera also revealed a dominant class 1 RBD pAb specificity, identical to the published mAb P4A1. Of interest, P4A1 uses IGHV3–53, a germline family frequently encoded by RBD antibodies from SARS-CoV-2-infected and mRNA-vaccinated donors ^26,70,71^.

In addition to RBM responses, both vaccines induced strong NTD responses, signifying the importance of understanding NTD antigenic sites targeted by different vaccines. Notably, a larger diversity of NTD pAb specificities were observed in donors vaccinated with NVX-CoV2373 compared to mRNA-1273 vaccinees. We have previously shown that the Spike NTD in the NVX- CoV2373 formulation exists in a more stable conformation compared to other published structures of Spike ectodomains ^48^, which may enable the induction of a larger diversity of NTD responses compared to mRNA vaccines. The NTD is a key target of infection-induced immunity and is under constant immune pressure as evidenced by numerous mutations, including insertions and deletions, observed in the NTD of VOCs^17,25,72^. These occur exclusively in the loops N1–5, and predominantly in N3, N5, and N1 that comprise the NTD supersite. In contrast to the RBD, which is the main target for neutralizing antibodies^73^, only a small fraction of antibodies to NTD are neutralizing, which primarily target the NTD supersite ^32–34^. Here, we show that both mRNA-1273 and NVX-CoV2373 elicit a dominant response to the supersite, which are highly specific to the ancestral Spike, exhibiting no cross-reactivity to other variants. These include 4A8-like antibodies commonly encoded by the public clonotype heavy chain VH1-24 gene that specifically target the N3 and N5 loops in a distinct manner ^32,34^. More recently, other epitopes on NTD, targeted by mAbs C1717, C1520, C1791, 5-7, and CoV2-3434, have also been shown to possess neutralizing ability while also exhibiting higher levels of sequence conservation ^56,74,75^. However, immunization with either vaccine did not elicit detectable levels of these antibody specificities. Of interest, certain NTD antibodies isolated from SARS-CoV-2-infected donors have also been shown to enhance infection by augmenting an RBD-open conformation or by other Fc-independent mechanisms ^55,63^. These antibodies mainly target the membrane-proximal face of NTD, a specificity that is commonly observed in both mRNA-1273 and NVX-CoV2373 immunized NHPs and clinical trial donors. Our findings not only underscore the need for a more detailed evaluation of the functional profiles of NTD-specific pAb responses elicited by various vaccines but also to develop strategies to shift response away from these highly variable immunodominant epitopes.

## Resource availability Lead contact

Reagent and resource requests should be addressed to and will be fulfilled by the lead contact, Andrew B. Ward

## Materials availability

Stabilized spike constructs are available upon request

## Data availability

All data needed to evaluate the conclusions in the paper are present in the paper and/or the Supplementary Materials. Cryo-EM maps have been deposited at the Electron Microscopy Data Bank (EMDB) with accession codes EMD-48188 (ModWu-RBD1), EMD-48189 (ModWu-RBD2), EMD-48190 (ModWu-NTD1), EMD-48192 (ModWu-NTD2), EMD-48193 (ModWu-NTD3), EMD-48194 (ModWu-NTD4), EMD-48195 (ModWu-NTD5), EMD-48196 (ModWu-NTD6), EMD-48197 (ModDe-RBD1), EMD-48198 (ModDe-NTD1), EMD-48199 (NVX-NHPWu1), EMD-48200 (NVX-NHPWu2), EMD-48201 (NVX-NHPWu3), EMD-48202 (NVX-NHPWu4), EMD-48203 (NVX-NHPWu5), EMD-48204 (NVX-NHPWu6), EMD-48205 (NVX-NHPWu7), EMD-48206 (NVX-NHPSA1), EMD-48207 (NVX-NHPSA2), EMD-48208 (NVX-NHPSA3), EMD-48209 (NVX-NHPSA4), EMD-48210 (NVX-NHPSA5), EMD-48211 (NVX-NHPSA6), EMD-48212 (NVX-NHPSA7) and EMD-48213 (NVX-NHPSA8).

## Acknowledgements

We thank Hannah L. Turner and Charles A. Bowman for their help with electron microscopy, data acquisition and data processing. We thank Lauren Holden for her assistance with the manuscript. We thank Ricardo Carrion, Jr. and Yenni Goez-Gazi at Texas Biomedical Research Institute for conducting the Novavax NHP experiment. We acknowledge Lynn Green and Emily Hoopii-Taylor on the Novavax Clinical Immunology Sample Management Team for preparing and coordinating shipment of human samples. mRNA-1273 work has been funded by the Intramural Research Program of the VRC, NIAID, NIH and in part with Federal funds from the Department of Health and Human Services, Office of the Assistant Secretary for Preparedness and Response, Biomedical Advanced Research and Development Authority, under Contract 75A50120C00034. KSC’s research fellowship was partially funded by the Undergraduate Scholarship Program, Office of Intramural Training and Education, Office of the Director, NIH.

## Author contributions

SB, AMJ and ABW conceptualized the work, designed experiments, interpreted results and wrote the manuscript. SB, AMJ, JC, MLF-Q, JLT, STR, BN, LMS and ATdlP acquired the data. AR, MG-X, NP and GS collected, handled and coordinated shipment of serum samples from Novavax NHP studies and clinical trials. KSC, RAS and BSG designed and conceptualized NHP studies with mRNA-1273 immunizations. BG, RD, and DKE collected, handled and coordinated shipment of serum samples from mRNA-1273 clinical trials.

## MATERIALS AND METHODS

### Expression and purification of recombinant Spike proteins

For the ancestral SARS-CoV-2 Spike proteins, we synthesized a base construct with residues 1 to 1208 from the Wuhan-Hu-1 strain (GenBank: QHD43416.1) with 6 stabilizing proline (HP) substitutions at positions 817, 892, 899, 942, 986, and 987 as described previously ^47^. The Spike ectodomain construct contains a C-terminal T4 fibritin trimerization domain, an HRV3C cleavage site, and a Twin-strep-tag for purification. The S1/S2 furin cleavage site was either modified to 682-GSAS-685 (GSAS) or kept as the WT sequence (RRAR). In addition to these, D614G mutation along with other interprotomer disulfide bonds, Mut2 (S383C, D985C), Mut4 (A570C, L966C), and Mut7 (T883C, V705C), were introduced as needed. A total of twelve different Wuhan S proteins were used for EM with ancestral Spike, including SARS-CoV-2-HP-GSAS; SARS-CoV-2-HP-GSAS-Mut2; SARS-CoV-2-HP-GSAS-Mut4; SARS-CoV-2-HP-GSAS-Mut7; SARS-CoV-2-HP-RRAR; SARS-CoV-2-HP-Mut2-RRAR; SARS-CoV-2-HP-Mut4-RRAR; SARS-CoV-2-HP-Mut7-RRAR; SARS-CoV-2-HP-GSAS-D614G; SARS-CoV-2-HP-GSAS- Mut2-D614G; SARS-CoV-2-HP-GSAS-Mut4-D614G; and SARS-CoV-2-HP-GSAS-Mut7- D614G.

The variant Spikes were generated from the HP-GSAS or HP-GSAS-Mut7 backbone with introduction of additional mutations; Alpha B.1.1.7_SARS-CoV-2-HP-GSAS-Mut7-D614G (69- 70 deletion, Y144 deletion, N501Y, A570D, D614G, P681H, T716I, S982A, and D1118H), Beta B.1.351_SARS-CoV-2-HP-GSAS-Mut7-D614G (D80A, D215G, K417T, E484K, N501Y, D614G, and A701V), Delta B.1.617.2_SARS-CoV-2-HP-GSAS-Mut7 (T19R, G142D, 156-157 deletion, R158G, L452R, T478K, D614G, P681R, and D950N) and Omicron B.1.1.529_SARS-CoV-2-HP-GSAS-Mut7 (A67V, 69-70 deletion, T95I, 142-144 deletion, Y145D, N211 deletion, 214 insertion, G339D, S371L, S373P, S375F, K417N, N440K, G446S, S477N, T478K, E484A, Q493R, G496S, Q498R, N501Y, and Y505H). S protein expression and purification were carried out as in previously published ^1^. Briefly, FreeStyle 293-F cells were transected with the Spike plasmid of interest and cultures were harvested at 6 days post-transfection. Spikes were purified using Strep-Tactin^®^ XT 4FLOW high-capacity columns (IBA Lifesciences) followed by SEC purification. Protein fractions corresponding to the trimeric Spike proteins were collected, concentrated, and assessed for quality by negative stain EM.

### mRNA-1273 NHP studies

Animal experiments were carried out in compliance with all pertinent US National Institutes of Health regulations and approval from the Animal Care and Use Committee of the Vaccine Research Center and Bioqual, Inc. (Rockville, MD). Studies were conducted at Bioqual, Inc. Twelve female and 12 male, 3-5 years old, Indian-origin rhesus macaques were immunized IM at week 0 and week 4 with 100 µg mRNA-1273 in 1 mL of 1X PBS into the right hindleg. At week 6, sera were collected for these analyses.

### Novavax NHP studies

The vaccination of NHPs with NVX-CoV2373 (Wuhan-Hu-1 spike protein) and rS-Beta (Beta variant spike protein) was performed at the Texas Biomedical Research Institute (San Antonio, TX). Female and male rhesus macaques (*Macaca mulatta*), 2-10 years old and weighing 3.1-11.3 kilograms at study initiation, were obtained from SNPRC specific pathogen free (SPF) colony and/or Envigo (Alice, TX). Animals were maintained at Texas Biomedical Research Institute for the entire in-life portion of the study and were treated according to Animal Welfare Act regulations and the Guide for the Care and Use of Laboratory Animals (2011). Animals were immunized via intramuscular (IM) injection with 3 μg of SARS-CoV-2 rS constructs (NVX-CoV2373 or rS-Beta) with 50 μg Matrix-M adjuvant administered as a prime/boost regimen, on Days 0 and 21, either as homologous (NVX-CoV2373/NVX-CoV2373 and rS-Beta/rS-Beta) or heterologous prime boost (NVX-CoV2373/rS-Beta) with 50 μg Matrix-M adjuvant. Blood was collected before the first immunization on Day 0 and Day 35 (2 weeks following the second immunization).

### Clinical trial studies

Novavax’s clinical samples were collected during Trial 2019nCoV-101 (ClinicalTrials.gov ID NCT04368988), a two-part, randomized, observer-blinded, placebo-controlled, Phase 1/2 study designed to evaluate the safety and immunogenicity of SARS-CoV-2 rS with and without Matrix-M adjuvant ^50,76^ . Participants were immunized intramuscularly with 5 μg NVX-CoV2373 with 50 μg Matrix-M adjuvant on Days 0 and 21. Sera collected on Day 21, 49, and 105 were analyzed in the current study.

Moderna’s clinical samples were collected during trial mRNA-1273-P101 (Clinicaltrials.gov ID NCT04283461), a Phase I, Open-Label, Dose-Ranging Study of the Safety and Immunogenicity of 2019-nCoV Vaccine (mRNA-1273) in Healthy Adults. Participants received an intramuscular (IM) injection (0.5 milliliters [mL]) of mRNA-1273 on Days 1 and 29. Sera collected on D29, and D43 was analyzed for the current study.

### Serum IgG Isolation and Digestion

Serum IgG isolation and digestion was performed as previously described^77^. Human serum samples were heat inactivated at 56°C for 1 h and then incubated with an equal volume of Protein G resin (GE Healthcare) for 3 nights at 4°C on a rocking platform. NHP serum samples were heat inactivated at 56°C for 1 h and then incubated with an equal volume of Protein A resin (GE Healthcare) for 3 nights at 4°C on a rocking platform. For both sample types, 1 x PBS pH 7.4 was used as a diluent to promote resin movement. Following incubation, resin was washed three times with 1 x PBS pH 7.4. IgG was eluted off resin using 5 resin-column volumes of 0.1 M Glycine pH 2.0, and the eluate was neutralized with 4 resin-column volumes of 1 M Tris pH 8.0. Neutralized eluate was filtered using a 0.22-μm Steriflip^®^ filtration unit (MilliporeSigma) and buffer exchanged to 1 x PBS pH 7.4 using either a 100kDa Amicon^®^ centrifugal filter unit (MilliporeSigma) or a 100kDa Pierce protein concentrator unit (Thermo Scientific). For Fab preparation, IgGs were incubated for 4–5 h at 37°C in digestion buffer (4% (w/w of IgG) activated Papain from papaya latex (Sigma Aldrich), 10 x L-Cysteine, 10 x Tris/EDTA pH8, ddH_2_0). Digestion reactions were quenched using 3x volume activated Papain of 0.5 M Iodoacetamide. Digested IgGs, now Fab/Fc, were buffer exchanged to 1 x TBS pH 7.4 using either 10kDa Amicon^®^ centrifugal filter units (MilliporeSigma) or 10kDa Pierce™ protein concentrator units (Thermo Scientific). The Fab/Fc was purified over a Superose 200 increase 10/300 column (Cytiva) to remove any undigested IgG, and the isolated Fab/Fc peak was concentrated using a 10kDa Amicon^®^ centrifugal filter unit (MilliporeSigma).

### Preparation of Polyclonal Fab-S Protein Complexes for Negative Stain-EM

For NHP Fab-Spike protein complexes, 250 μg of purified polyclonal Fab was complexed with 15 μg of a mix of stabilized Spike proteins (Moderna complexes: SARS-CoV-2-6P-RRAR, SARS-CoV-2-6P-Mut2-RRAR, SARS-CoV-2-6P-Mut4-RRAR, SARS-CoV-2-6P-Mut7-RRAR, SARS-CoV-2-6P-D614G, SARS-CoV-2-6P-Mut2-D614G, SARS-CoV-2-6P-Mut4-D614G, SARS-CoV-2-6P-Mut7-D614G. Novavax complexes: SARS-CoV-2-6P-D614G, SARS-CoV-2-6P- Mut2-D614G, SARS-CoV-2-6P-Mut4-D614G, SARS-CoV-2-6P-Mut7-D614G, SARS-CoV-2-6P-Mut7) or a variant Spike protein. Moderna NHP Fab-Spike protein complexes were incubated for 7–15 min at room temperature and crosslinked according to published methods^53^. Novavax NHP samples were complexed overnight at room temperature. For both Moderna and Novavax clinical trial samples, 1 mg of purified polyclonal Fab was mixed with 15 μg of a mix of stabilized Spike proteins (SARS-CoV-2-6P-D614G, SARS-CoV-2-6P-Mut2-D614G, SARS-CoV-2-6P-Mut4-D614G, SARS-CoV-2-6P-Mut7-D614G, SARS-CoV-2-6P-Mut7) or a variant Spike protein. These complexes were incubated for 2 h at room temperature and crosslinked according to published methods. Complexes were purified over a Superose 6 increase 10/300 column (Cytiva) using UV 215 absorbance on an ӒKTA pure™ system (Cytiva). Protein fractions corresponding to the Spike protein were collected and concentrated using a 0.5-mL capacity 10kDa Amicon^®^ Ultra Centrifugal Filter Unit (MilliporeSigma).

### Negative Stain Electron Microscopy

Negative stain electron microscopy was performed as previously described (Ref). In brief, samples were diluted to around 0.03 mg/mL in 1 x TBS at pH 7.4 and applied onto carbon coated 400- mesh copper grids. Grids were stained with either 2% Nano-W™ (Nanoprobes) or 2% (w/v) uranyl formate. Nano-W grids were stained for 7 s then again for 14 s. Uranyl formate grids were stained for 90 s. Leginon software^78^ was used for automated data collection on various microscopes including: a 120keV FEI Tecnai Spirit electron microscope with a Thermo Fisher Eagle 4k x 4k CCD camera at 52,000x magnification, -1.50 μm nominal defocus, and 2.063 Å pixel size; a 200keV FEI Tecnai F20 electron microscope with a T-VIPS TemCam F416 CMOS camera at 62,000x magnification, -1.50 μm nominal defocus, and 1.677 Å pixel size; and a FEI Talos F200X G2 electron microscope with a Thermo Fisher CETA 4K CMOS camera at either a 73,000x magnification and 1.981 Å pixel size or a 92,000x magnification and 1.572 Å pixel size. DoGpicker^3^ was used to pick particles via Appion^4^, and particles were processed in RELION 3.0^5^. An initial model was generated from a published SARS-CoV-2 S protein structure (PDB: 6VYB^6^) and used during data processing. Segmentation for figures was performed using either UCSF Chimera^7^ or UCSF ChimeraX^8^.

### Cryo-EM sample preparation

Cryo-EM complexes for NVX-CoV2373-immunized NHPs were prepared by pooling equal amounts of polyclonal Fabs from four animals and complexing 1 mg of the pooled Fab with 80 μgs of Wuhan SARS-CoV-2-HP-D614G-Mut7. Immune complexes for heterologous prime boosting with NVX-CoV2373/rS-B.1.35 were prepared by complexing 1 mg of polyclonal Fab from an individual representative NHP with 80 μg of Wuhan SARS-CoV-2-HP-D614G-Mut7 spike. Complexes were incubated overnight at room temperature and purified as described for ns- EMPEM complexes. For grid preparation, 3.5 µl of purified NHP polyclonal complexes (concentrations of ∼0.4 µg/mL) were mixed with 0.5 µl of 0.04 mM Lauryl maltose neopentyl glycol (LMNG) solution immediately before sample deposition onto QUANTIFOIL^®^ 2/1 400 mesh grids (EMS), which were plasma cleaned for 10 s using Gatan Solarus 950 Plasma system (Ar/O_2_ gas mixture). Following sample application, grids were blotted for 3.5 s before being plunged into liquid nitrogen-cooled liquid ethane using a Vitrobot Mark IV (Thermo Fisher Scientific).

Mod-D43 Fab-Spike protein complexes for Moderna clinical trial samples were made by mixing 4 mg of purified polyclonal Fabs (pooled from four donors) with 80 μg of either the Wuhan SARS-CoV-2-HP-D614G-Mut7 S protein or the Delta SARS-CoV-2-HP-D614G-Mut7 Spike. Complexes were incubated for 2 h at room temperature, crosslinked and purified as described previously. Grids were prepared by mixing 3.5 µl of purified polyclonal complexes (concentrations of ∼0.5 µg/mL) with either 0.5 µl of 0.04 mM lauryl maltose neopentyl glycol (LMNG) or 0.5 µl of 0.48 mM *n*-dodecyl-B-D-maltopyranoside (DDM) solutions immediately before deposition onto plasma-cleaned Quantifoil 2/1 grids (EMS). Grids were either blotted for 3 or 4 s before being plunged into liquid nitrogen-cooled liquid ethane using a Vitrobot mark IV (Thermo Fisher Scientific).

### Cryo-EM data collection and processing

For the NHP polyclonal complexes, micrographs were collected on a FEI Titan Krios operating at 300 keV mounted with a Gatan K2 direct-electron detector through Leginon software. The collection parameters are described in Table **S1**. MotionCor2^79^ was used for alignment and dose weighting of the frames. Initial processing was performed on CryoSPARC^80^, including CTF estimations (GCTF)^81^, micrograph curation based on CTF resolution estimates (cutoff 5 Å), template-based particle picking, particle extraction, and iterative rounds of 2D classification to select for intact Spike trimers. The clean particle stacks were then transferred to Relion^82^ for 3D refinement and different antibody classes were sorted using the focused classification protocol described previously.

Micrographs for Mod-D42 Fabs complexed with both Wuhan and Delta Spikes were collected at PNCC on a FEI Titan Krios operating at 300 keV mounted with a Gatan K3. Frames were aligned using Relion MotionCor2 and micrographs were transferred to Cryosparc for initial processing as previously described for the NHP datasets. The clean particle stacks were then transferred to Relion for 3D refinement, C3 symmetry expansion, followed by focused classification to sort the different antibody classes as described previously.

The collection and final refinement parameters are provided in Tables **S1–S4** and a schematic representation of processing workflows for each cryo-EMPEM dataset are shown in Figures **S3, S4, S7, and S8**.

### Molecular dynamics simulations

As starting structure for our simulations, we used the available cryo-EM structure of the full-length SARS-CoV-2 prefusion spike trimer (PDB accession code: 7JJI), adding the HexaPro mutations reported to stabilize the Spike trimer using MOE (*Molecular Operating Environment (MOE)*, 2022.02 Chemical Computing Group ULC, 910-1010 Sherbrooke St. W., Montreal, QC H3A 2R7, **2024**.) We performed three repetitions of 600 ns of classical molecular dynamics simulations using the AMBER 22 simulation software package which contains the pmemd.cuda module (D.A. Case, H.M. Aktulga, K. Belfon, I.Y. Ben-Shalom, J.T. Berryman, S.R. Brozell, D.S. Cerutti, T.E. Cheatham, III, G.A. Cisneros, V.W.D. Cruzeiro, T.A. Darden, R.E. Duke, G. Giambasu, M.K. Gilson, H. Gohlke, A.W. Goetz, R. Harris, S. Izadi, S.A. Izmailov, K. Kasavajhala, M.C. Kaymak, E. King, A. Kovalenko, T. Kurtzman, T.S. Lee, S. LeGrand, P. Li, C. Lin, J. Liu, T. Luchko, R. Luo, M. Machado, V. Man, M. Manathunga, K.M. Merz, Y. Miao, O. Mikhailovskii, G. Monard, H. Nguyen, K.A. O’Hearn, A. Onufriev, F. Pan, S. Pantano, R. Qi, A. Rahnamoun, D.R. Roe, A. Roitberg, C. Sagui, S. Schott-Verdugo, A. Shajan, J. Shen, C.L. Simmerling, N.R. Skrynnikov, J. Smith, J. Swails, R.C. Walker, J. Wang, J. Wang, H. Wei, R.M. Wolf, X. Wu, Y. Xiong, Y. Xue, D.M. York, S. Zhao, and P.A. Kollman (2022), Amber 2022, University of California, San Francisco.). The structure was prepared using CHARMM-GUI^83,84^. The structure models were placed into cubic water boxes of TIP3P water molecules with a minimum wall distance to the protein of 12 Å^85^. Parameters for all simulations were derived from the AMBER force field 14SB^86^. To neutralize the charges, we used uniform background charges^87^. The protein was carefully equilibrated using a multistep equilibration protocol^88^.

Bonds involving hydrogen atoms were restrained using the SHAKE algorithm, allowing a timestep of 2.0 femtoseconds. The systems’ pressure was maintained at 1 bar by a Monte Carlo barostat with one volume change attempt per 100 steps. The Langevin Thermostat was utilized to keep the temperature at 300K during the simulations.

To characterize the different observed NTD conformations, we performed a cluster analysis implemented in cpptraj aligning on the whole spike protein and clustering solely on the NTD Cα-atoms using a distance cut-off criterion of 2.5 Å. Furthermore, we used an in-house python Script to identify the residues that are mainly solvent exposed in different conformational states, as an additional criterion for the identified residues that contribute to antibody recognition.

