## Supplemental Material for "Structural serology of polyclonal antibody responses to mRNA-1273 and NVX-CoV2373 COVID-19 vaccines"

[1]Dept. of Integrative Structural and Computational Biology, The Scripps Research Institute; La Jolla, CA, 92037, USA
[2]Novavax, Inc; 21 Firstfield Road, Gaithersburg, MD, 20878, USA
[3]Moderna, Inc; Cambridge, MA, 02142, USA
[4]Vaccine Research Center; National Institutes of Allergy and Infectious Diseases; National Institutes of Health; Bethesda, Maryland, 20892, USA
[5]Current affiliation: Department of Immunology and Infectious Diseases; Harvard T.H. Chan School of Public Health; Boston, Massachusetts, 02115, USA
[6]Current affiliation: Howard Hughes Medical Institute; Chevy Chase, Maryland, 20815, USA
[7]Current affiliation: Department of Microbiology, Biochemistry & Immunology; Morehouse School of Medicine; Atlanta, Georgia, 30310, USA
[#]These authors contributed equally
***Correspondence:**
Andrew B. Ward

#### Figures

A.

| Animal ID | D35 Anti-rS (EC50) | D35 Plaque Reduction Neutralization Titers (PRNT50) USA-WA1/2020 |
|---|---|---|
| NVXWu-NHP1 | 62222 | 1673.29 |
| NVXWu-NHP2 | 168573 | 3568.71 |
| NVXWu-NHP3 | 33412 | 1074.24 |
| NVXWu-NHP4 | 327608 | 4371.4 |

B.

| Animal ID | ID50 Titer (Week6) |
|---|---|
| Mod-NHP1 | 3323 |
| Mod-NHP2 | 4036 |
| Mod-NHP3 | 7299 |
| Mod-NHP4 | 8609 |

C.

| Pooled Groups | Animal ID | D35 Anti-rS Wuhan (EC50) | D35 Anti-rS SA (EC50) | D35 SARS-CoV-2 Plaque Reduction Neutralization Titers (PRNT50) | | | |
|---|---|---|---|---|---|---|---|
| | | | | USA-WA1/2020 | Beta B.1.351 | Alpha B.1.1.7 | Delta B.1.617.1 |
| Homologous NVX-CoV2373 prime-boost | NVXWu-NHP1 | 62222 | 40288 | 1673.29 | 650.25 | 4684.25 | 4926.17 |
| | NVXWu-NHP2 | 168573 | 97100 | 3568.71 | 2097.99 | 8608.69 | 5201.07 |
| | NVXWu-NHP3 | 33412 | 21991 | 1074.24 | 324.86 | 1010.4 | 1349.95 |
| | NVXWu-NHP4 | 327608 | 235987 | 4371.4 | 4802.9 | 11525.65 | 22316.01 |
| | NVXWu-NHP5 | 181272 | 121172 | 4478.01 | 3104.66 | 12750.32 | 17762.63 |
| Homologous rS-B.1.351 prime-boost | NVXSA-NHP1 | 24418 | 51650 | 433.96 | 2640.67 | 2262.67 | 993.9 |
| | NVXSA-NHP2 | 176122 | 309877 | 504.45 | 14752.85 | 1732.13 | 1168.25 |
| | NVXSA-NHP3 | 83828 | 172131 | 404.07 | 6187.01 | 2828.59 | 346.84 |
| | NVXSA-NHP4 | 31414 | 60845 | 303.1 | 4167.7 | 1278.43 | 614.93 |
| | NVXSA-NHP5 | 235781 | 448973 | 1318.11 | 22091.49 | 9236.44 | 4289.18 |
| Heterologous NVX-CoV2373/rS-B.1.351 prime-boost | NVXWu/SA-NHP1 | 83327 | 81397 | 1156.28 | 587 | 2066.27 | 517.68 |
| | NVXWu/SA-NHP2 | 123916 | 129238 | 1574.24 | 2642.11 | 4383.54 | 411.63 |
| | NVXWu/SA-NHP3 | 275603 | 234235 | 3295.05 | 4594.49 | 6069.98 | 845.41 |
| | NVXWu/SA-NHP4 | 183030 | 167035 | 3736.96 | 5152.95 | 7773.29 | 2720.84 |

**Figure S1. NHP serum antibody responses to NVX-CoV2373, NVX rS-Beta, and mRNA-1273 (A)** Day 35 serum binding titers to recombinant Wuhan Spike shown as half-maximal effective concentration ($EC_{50}$) and WA1/2020 virus inhibition titers indicated as half-maximal plaque reduction neutralization titers ($PRNT_{50}$) from NHPs immunized with NVX-CoV2373 prime and boost. **(B)** Week 6 serum neutralizing titers shown as 50% inhibitory dose ($ID_{50}$) from NHPs immunized with mRNA-1273 prime-boost regimen. **(C)** Day 35 serum binding titers to recombinant Spikes (rS) from Wuhan and Beta (SA) strains shown as half-maximal effective concentrations ($EC_{50}$) and virus neutralization titers against WA1/2020, Beta B.1.351, Alpha B.1.1.7, and Delta B.1.617.1 viruses indicated as half-maximal plaque reduction neutralization titers ($PRNT_{50}$) from NHPs immunized with either homologous NVX-CoV2373 or rS-Beta prime and boost series or heterologous NVX-CoV2373 prime and rS-Beta boost. Samples highlighted in grey are the same samples as in (A). Sample highlighted in yellow was used for cryo-EM studies.

| | | |
|---|---|---|
| NVX-CoV2373 | NVX-Wu NHP1 + Wu Spike Mix | 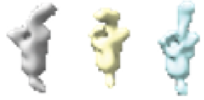 |
| | NVX-Wu NHP2 + Wu Spike Mix | 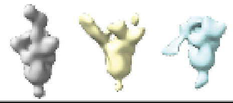 |
| | NVX-Wu NHP3 + Wu Spike Mix | 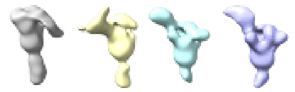 |
| | NVX-Wu NHP4 + Wu Spike Mix | 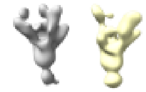 |
| mRNA-1273 | Mod NHP1 + Wu Spike Mix | 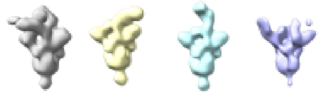 |
| | Mod NHP2 + Wu Spike Mix | 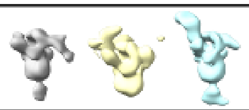 |
| | Mod NHP3 + Wu Spike Mix | 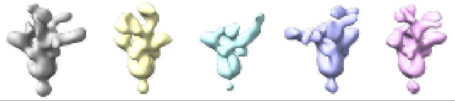 |
| | Mod NHP4 + Wu Spike Mix | 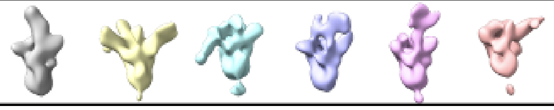 |
| NVX-CoV2373 | NVX-Donor2 (Day 21) + Wu Spike Mix | 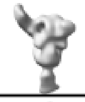 |
| | NVX-Donor3 (Day 21) + Wu Spike Mix | 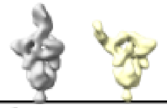 |
| | NVX-Donor4 (Day 21) + Wu Spike Mix | 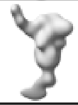 |
| | NVX-Donor1 (Day 49) + Wu Spike Mix | 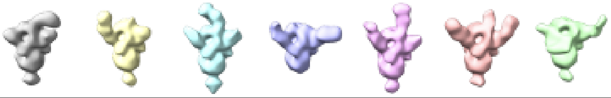 |
| | NVX-Donor2 (Day 49) + Wu Spike Mix | 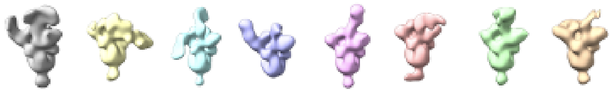 |
| | NVX-Donor3 (Day 49) + Wu Spike Mix | 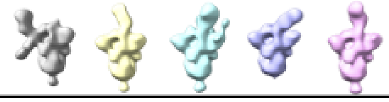 |
| | NVX-Donor4 (Day 49) + Wu Spike Mix | 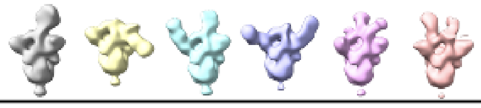 |
| | NVX-Donor5 (Day 49) + Wu Spike Mix | 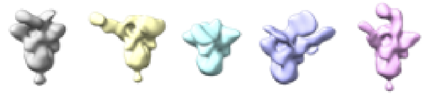 |

| | | |
|---|---|---|
| NVX-CoV2373 | NVX-Donor1 (Day 105) + Wu Spike Mix | 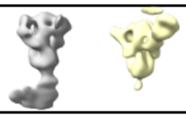 |
| | NVX-Donor2 (Day 105) + Wu Spike Mix | 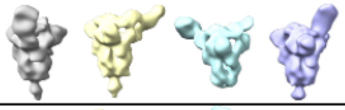 |
| | NVX-Donor3 (Day 105) + Wu Spike Mix | 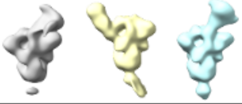 |
| | NVX-Donor4 (Day 105) + Wu Spike Mix | 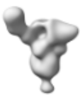 |
| mRNA-1273 | Mod-Donor1 (Day 29) + Wu Spike Mix | 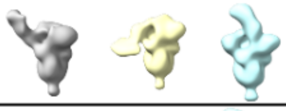 |
| | Mod-Donor2 (Day 29) + Wu Spike Mix | 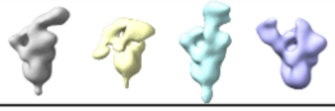 |
| | Mod-Donor3 (Day 29) + Wu Spike Mix | 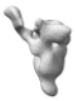 |
| | Mod-Donor4 (Day 29) + Wu Spike Mix | 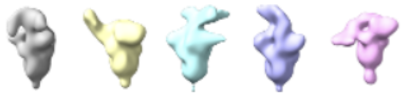 |
| | Mod-Donor1 (Day 43) + Wu Spike Mix | 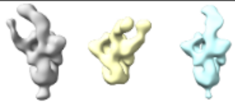 |
| | Mod-Donor2 (Day 43) + Wu Spike Mix | 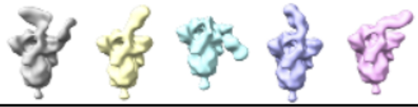 |
| | Mod-Donor3 (Day 43) + Wu Spike Mix | 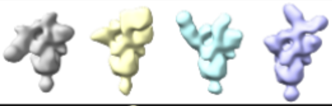 |
| | Mod-Donor4 (Day 43) + Wu Spike Mix | 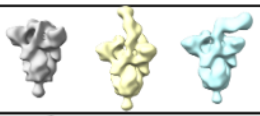 |
| NVX-CoV2373 | NVX-Wu NHP1 + Alpha Variant Spike | 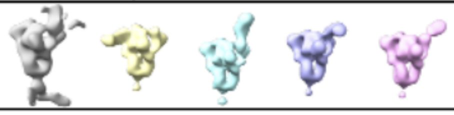 |
| | NVX-Wu NHP1 + Beta Variant Spike | 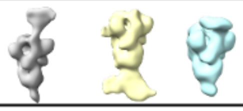 |

| | | |
|---|---|---|
| NVX-CoV2373 | NVX-Wu NHP2 + Alpha Variant Spike | 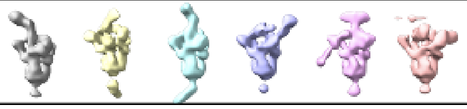 |
| | NVX-Wu NHP2 + Beta Variant Spike | 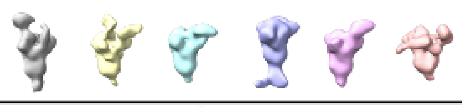 |
| | NVX-Wu NHP2 + Delta Variant Spike | 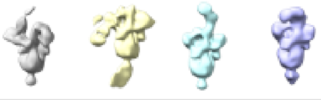 |
| | NVX-Wu NHP2 + Omicron Variant Spike | 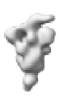 |
| mRNA-1273 | Mod NHP4 + Alpha Variant Spike | 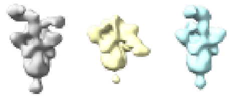 |
| | Mod NHP4 + Beta Variant Spike | 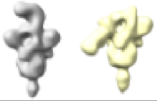 |
| | Mod NHP4 + Delta Variant Spike | 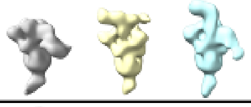 |
| | Mod NHP4 + Omicron Variant Spike | 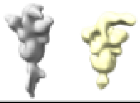 |
| | Mod NHP1 + Alpha Variant Spike | 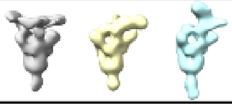 |
| | Mod NHP1 + Beta Variant Spike | 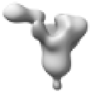 |
| | Mod NHP1 + Delta Variant Spike | 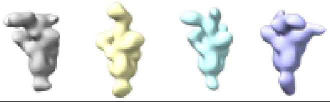 |
| | Mod NHP1 + Omicron Variant Spike | 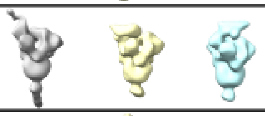 |
| NVX-CoV2373 | NVX- D49 Donor Pool + Ancestral Spike | 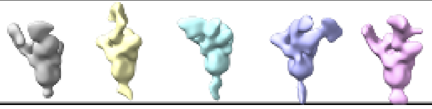 |
| | NVX- D49 Donor Pool + Beta Variant Spike | 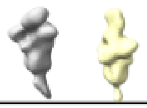 |
| | NVX- D49 Donor Pool + Delta Variant Spike | 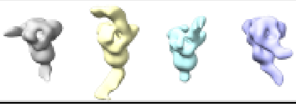 |
| | NVX- D49 Donor Pool + Omicron Variant Spike | 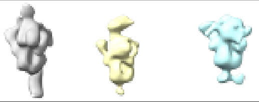 |

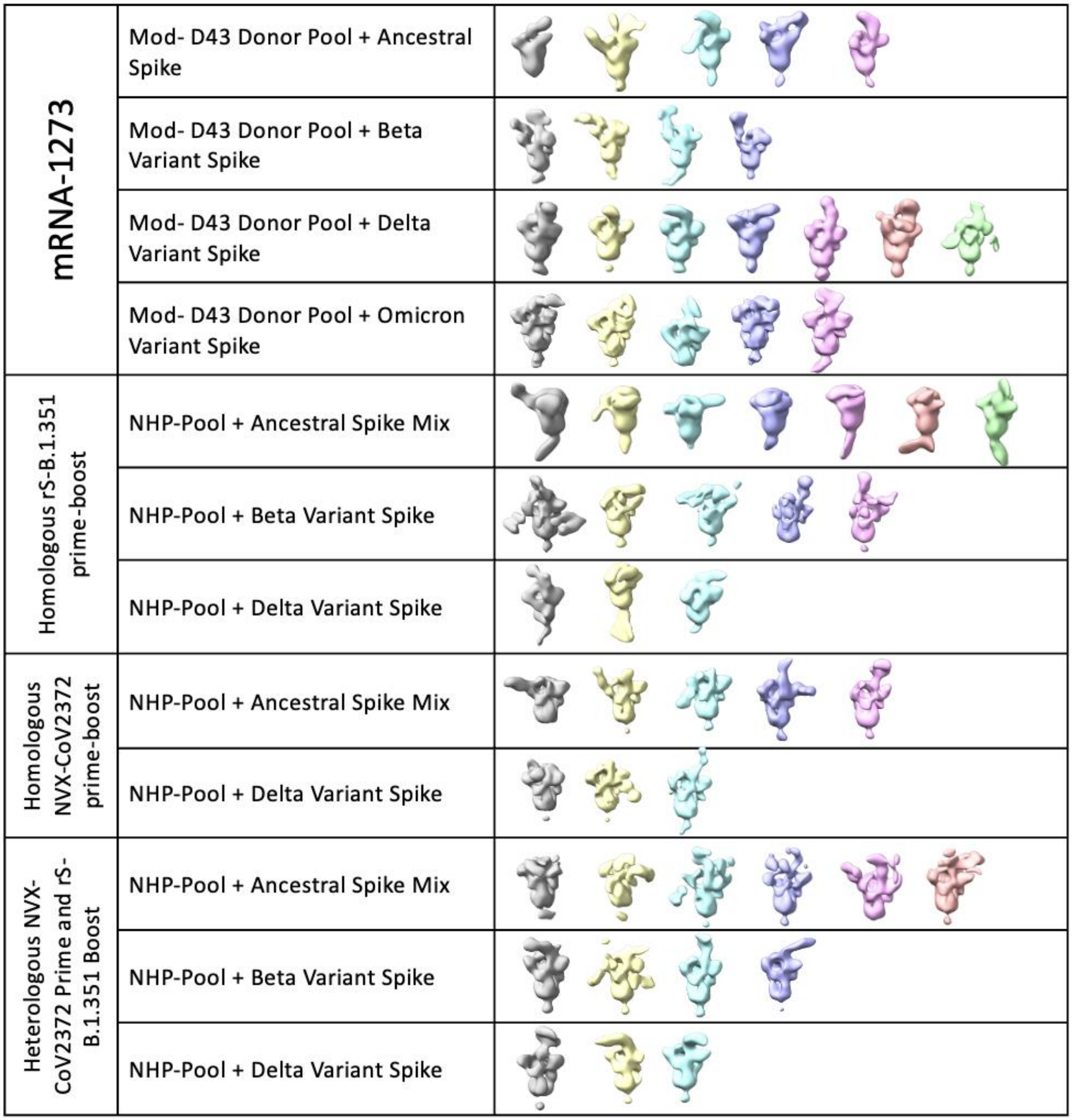


**Figure S2. Negative stain-EM 3D classes from each dataset used for generation of composite maps displaying all the specificities.**

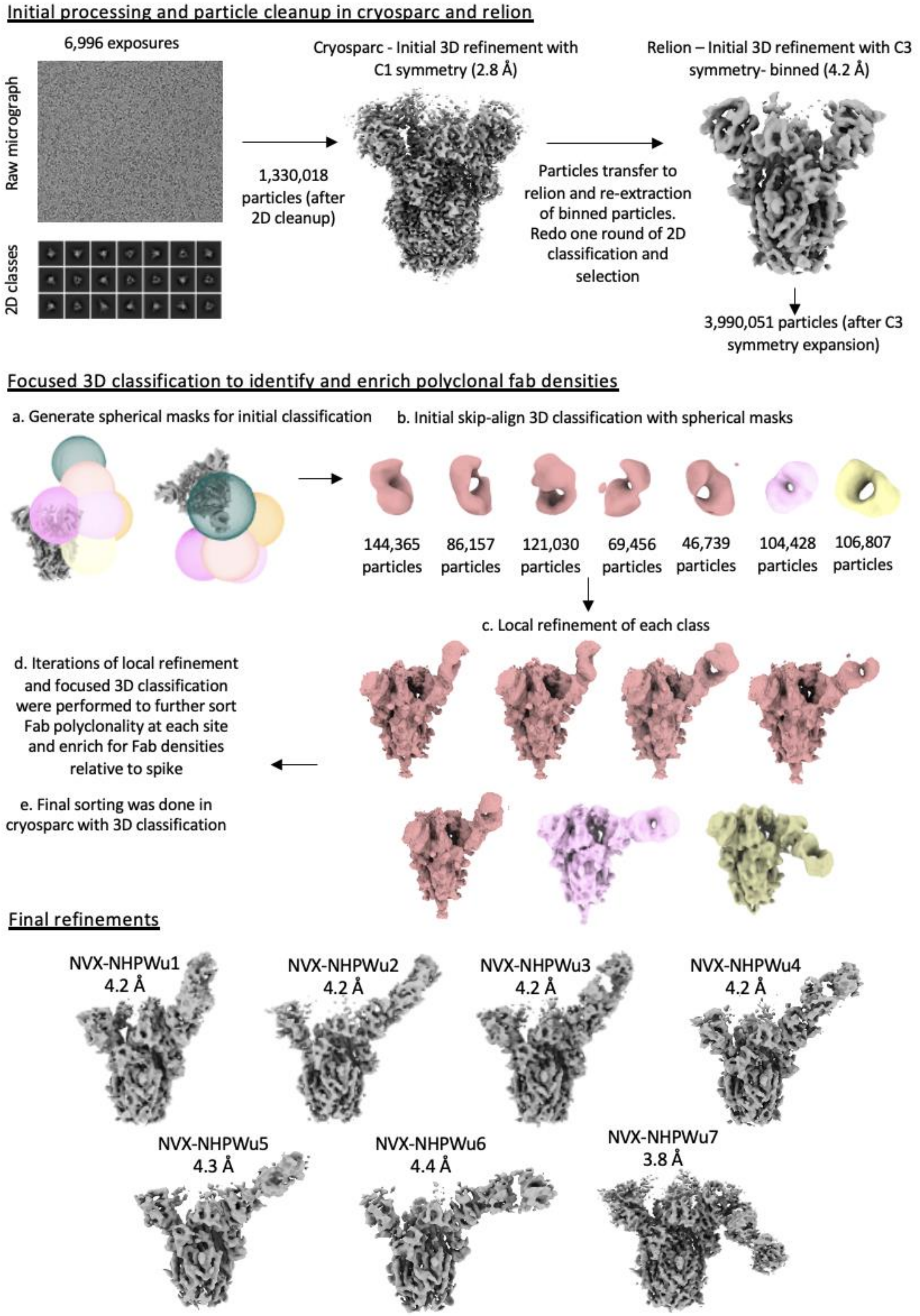


**Figure S3. Schematic representation of the cryo-EMPEM processing workflow for pooled NVX-CoV2373 prime-boosted NHP Fabs complexed with SARS-CoV-2-HP-GSAS-Mut7-**

**D614G Spike.** The focused classification approach used for generating Fab-Spike reconstructions is shown in steps.

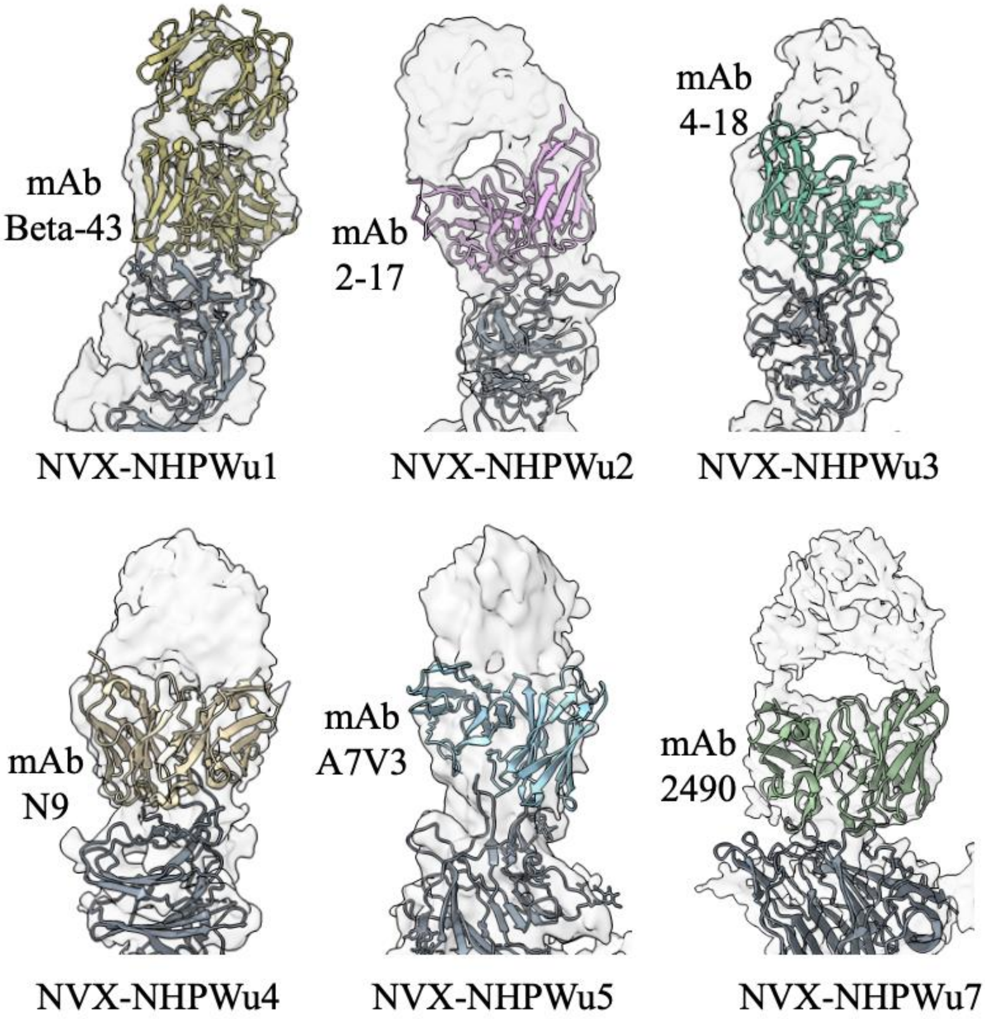


**Figure S4. Published monoclonal Spike-NTD antibodies resembling NVX-NHPWu pAbs.** Atomic models of NTD in complex with mAbs Beta-43 (PDB# 7Q91), 2-17 (PDB# 7LQW), 4-18 (PDB# 7L2E), N9 (PDB# 7E8F), A7V3 (PDB# 7SJ0), and 2490 (PDB# 7DZY) docked into cryo-EMPEM map densities (transparent) of pAbs NVX-NHPWu1, NVX-NHPWu2, NVX-NHPWu3, NVX-NHPWu4, NVX-NHPWu5, and NVX-NHPWu7, respectively.

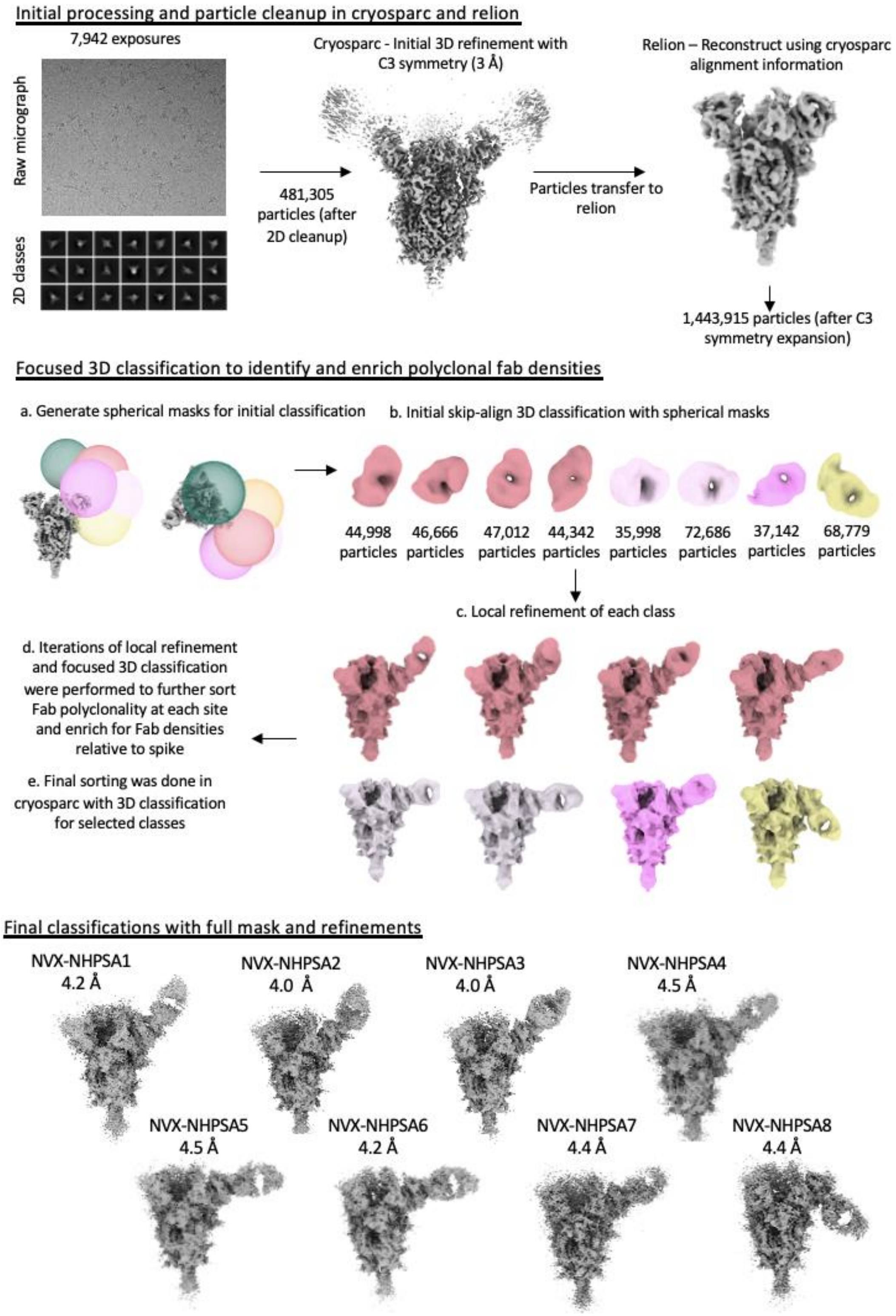

Initial processing and particle cleanup in cryosparc and relion
7,942 exposures
Raw micrograph
2D classes
481,305 particles (after 2D cleanup)
Cryosparc - Initial 3D refinement with C3 symmetry (3 Å)
Particles transfer to relion
Relion – Reconstruct using cryosparc alignment information
1,443,915 particles (after C3 symmetry expansion)
Focused 3D classification to identify and enrich polyclonal fab densities
a. Generate spherical masks for initial classification
b. Initial skip-align 3D classification with spherical masks
44,998 particles
46,666 particles
47,012 particles
44,342 particles
35,998 particles
72,686 particles
37,142 particles
68,779 particles
c. Local refinement of each class
d. Iterations of local refinement and focused 3D classification were performed to further sort Fab polyclonality at each site and enrich for Fab densities relative to spike
e. Final sorting was done in cryosparc with 3D classification for selected classes
Final classifications with full mask and refinements
NVX-NHPSA1 4.2 Å
NVX-NHPSA2 4.0 Å
NVX-NHPSA3 4.0 Å
NVX-NHPSA4 4.5 Å
NVX-NHPSA5 4.5 Å
NVX-NHPSA6 4.2 Å
NVX-NHPSA7 4.4 Å
NVX-NHPSA8 4.4 Å

**Figure S5. Schematic representation of the cryo-EMPEM processing workflow for SARS-CoV-2-HP-GSAS-Mut7-D614G Spike complexed with pAbs isolated from a single NHP immunized with NVX-CoV2373/rS-Beta prime-boost.**

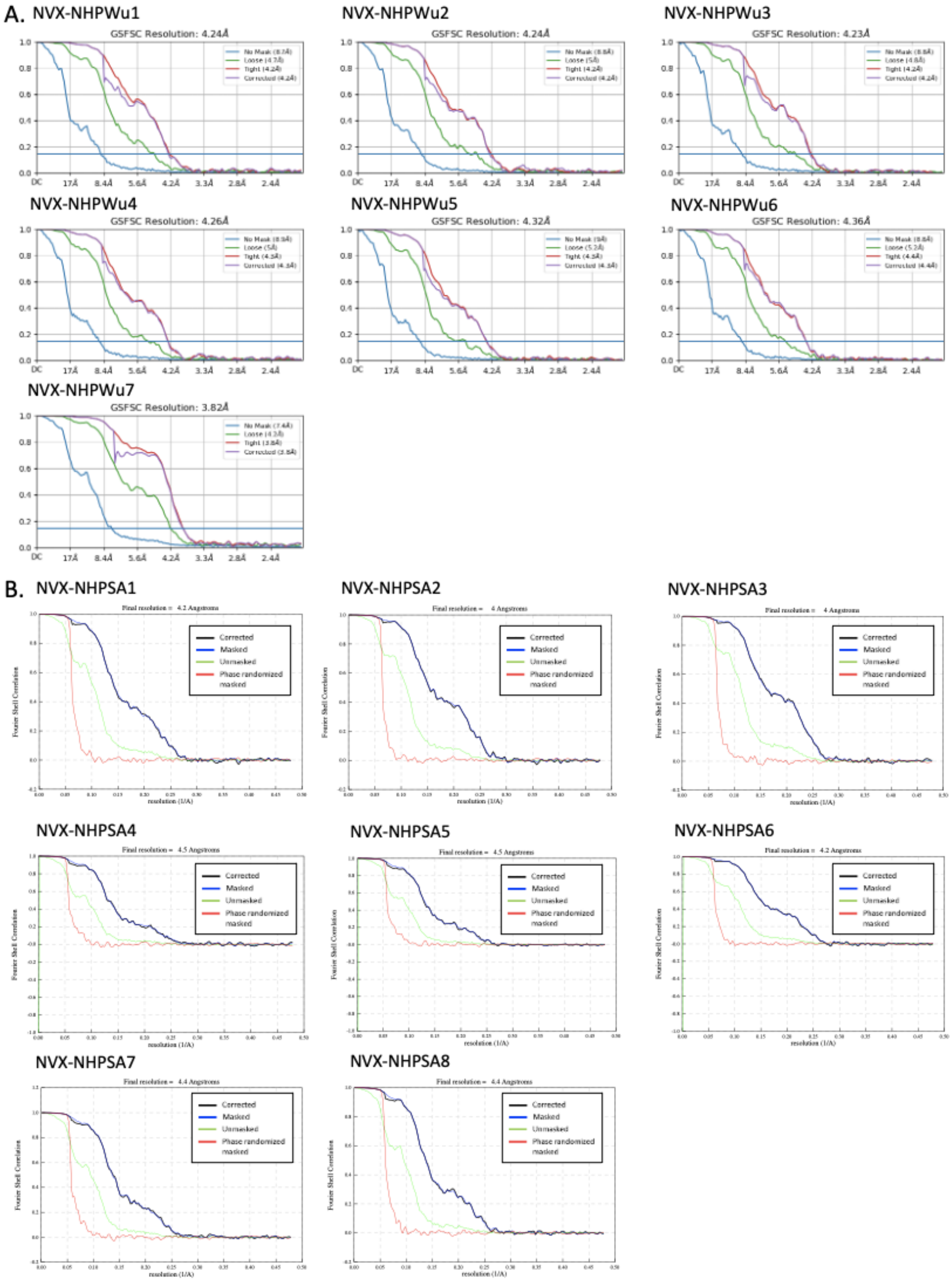


**Figure S6. FSC curves for SARS-CoV-2-HP-GSAS-Mut7-D614G Spike-Fab cryo-EMPEM reconstructions from NHPs that received (A) homologous NVX-CoV2373 or (B) heterologous NVX-CoV2373/rS-Beta prime-boost immunizations.**

A.

| Subject ID | Day | ELISA (IgG Titer) | ACE-2 Inhibition (Ab titer) | Micro neutralization (1/Dilution) |
|---|---|---|---|---|
| NVX-Donor1 | 21 | 3.62E+03 | 13.5 | 160 |
| NVX-Donor2 | 21 | 7.05E+03 | 5 | 80 |
| NVX-Donor3 | 21 | 2.22E+04 | 5 | 480 |
| NVX-Donor4 | 21 | 3.43E+03 | 12.9 | 240 |
| NVX-Donor5 | 21 | 4.06E+03 | 5 | 80 |
| NVX-Donor1 | 49 | 8.75E+04 | 173.1 | 7680 |
| NVX-Donor2 | 49 | 1.00E+05 | 288.2 | 10240 |
| NVX-Donor3 | 49 | 1.08E+05 | 184.6 | 5120 |
| NVX-Donor4 | 49 | 1.44E+05 | 435.3 | 15360 |
| NVX-Donor5 | 49 | 9.66E+04 | 344.9 | 5120 |
| NVX-Donor1 | 105 | 1.64E+04 | 33.8 | NA |
| NVX-Donor2 | 105 | 5.18E+04 | 64.4 | NA |
| NVX-Donor3 | 105 | 2.28E+04 | 14.8 | NA |
| NVX-Donor4 | 105 | 4.58E+04 | 155.2 | NA |
| NVX-Donor5 | 105 | 2.97E+04 | 77.5 | NA |

B.

| Subject ID | Day | Endpoint spike titer | Endpoint RBD titer | Pseudoneut titers |
|---|---|---|---|---|
| Mod-Donor1 | 29 | 1.06E+05 | 8.71E+04 | 21 |
| Mod-Donor2 | 29 | 9.54E+04 | 9.21E+04 | 14 |
| Mod-Donor3 | 29 | 9.23E+04 | 1.45E+05 | 22 |
| Mod-Donor4 | 29 | 1.69E+05 | 1.06E+05 | 69 |
| Mod-Donor1 | 43 | 5.57E+05 | 3.37E+05 | 271 |
| Mod-Donor2 | 43 | 7.08E+05 | 5.42E+05 | 361 |
| Mod-Donor3 | 43 | 7.11E+05 | 3.97E+05 | 320 |
| Mod-Donor4 | 43 | 8.28E+05 | 7.17E+05 | 697 |

**Figure S7. Clinical trial donor serum antibody responses to NVX-CoV2373 and mRNA-1273 at pre-boost and post-boost time points (A)** Day 21, day 49 and day 105 serum binding titers to recombinant Wuhan Spike shown as half-maximal effective concentration ($EC_{50}$), ACE2 inhibition titers and wild-type WA1/2020 microneutralization titers indicated as inhibitory concentration of >99% (MN $IC_{>99\%}$) from donors who received NVX-CoV2373 prime and boost. **(B)** Day 29 and day 43 serum binding titers to recombinant Wuhan Spike and recombinant RBD shown as half-maximal effective concentration ($EC_{50}$), and Wuhan pseudovirus neutralization titers from donors who received mRNA-1273 prime and boost.

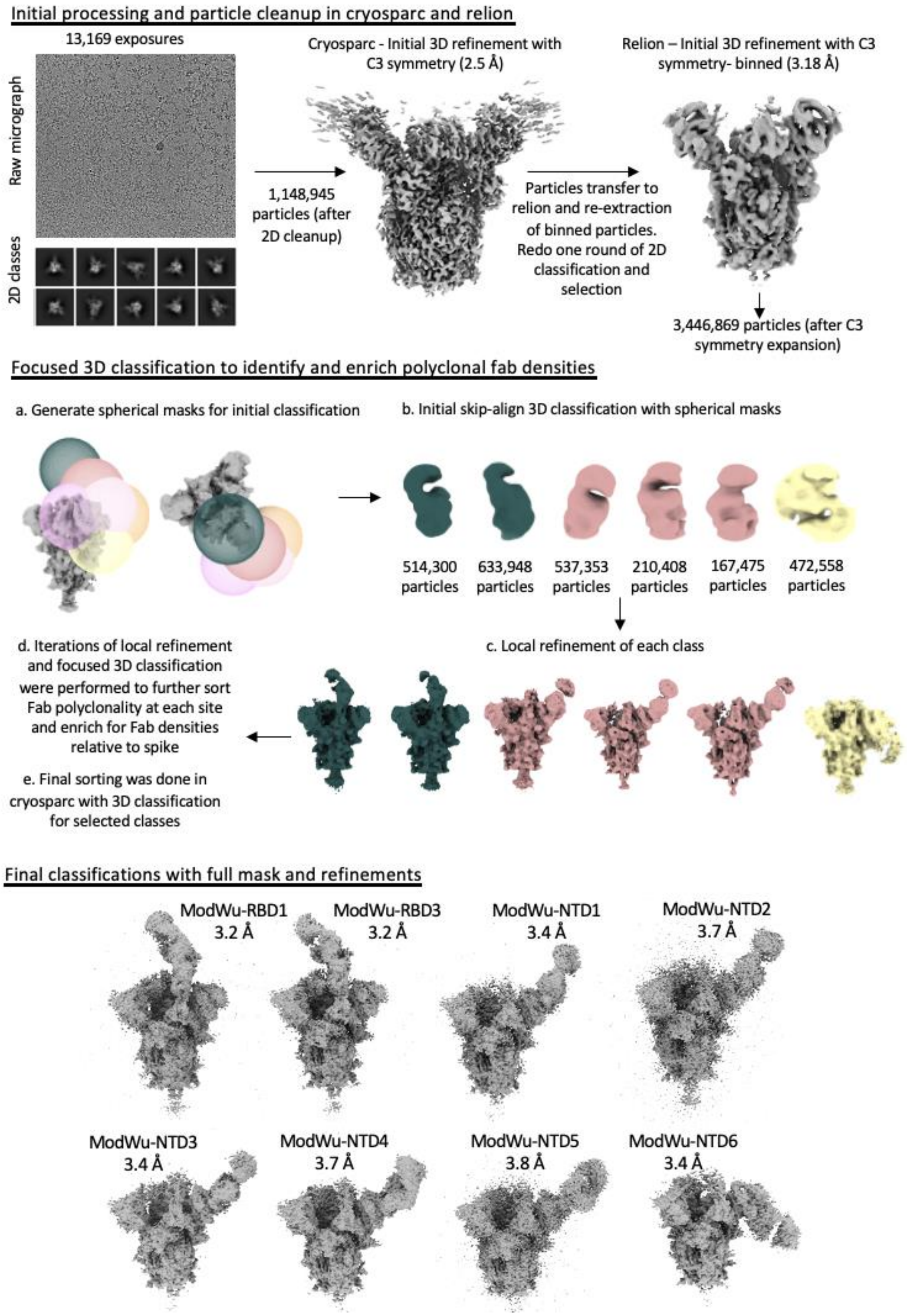
Initial processing and particle cleanup in cryosparc and relion
13,169 exposures
Raw micrograph
2D classes
1,148,945 particles (after 2D cleanup)
Cryosparc - Initial 3D refinement with C3 symmetry (2.5 Å)
Particles transfer to relion and re-extraction of binned particles. Redo one round of 2D classification and selection
Relion – Initial 3D refinement with C3 symmetry- binned (3.18 Å)
3,446,869 particles (after C3 symmetry expansion)
Focused 3D classification to identify and enrich polyclonal fab densities
a. Generate spherical masks for initial classification
b. Initial skip-align 3D classification with spherical masks
514,300 particles
633,948 particles
537,353 particles
210,408 particles
167,475 particles
472,558 particles
c. Local refinement of each class
d. Iterations of local refinement and focused 3D classification were performed to further sort Fab polyclonality at each site and enrich for Fab densities relative to spike
e. Final sorting was done in cryosparc with 3D classification for selected classes
Final classifications with full mask and refinements
ModWu-RBD1 3.2 Å
ModWu-RBD3 3.2 Å
ModWu-NTD1 3.4 Å
ModWu-NTD2 3.7 Å
ModWu-NTD3 3.4 Å
ModWu-NTD4 3.7 Å
ModWu-NTD5 3.8 Å
ModWu-NTD6 3.4 Å

**Figure S8. Schematic representation of the cryo-EMPEM processing workflow for pooled mRNA-1273 vaccinated donor serum pAbs (day 43) complexed with SARS-CoV-2-HP-GSAS-Mut7-D614G Spike.**

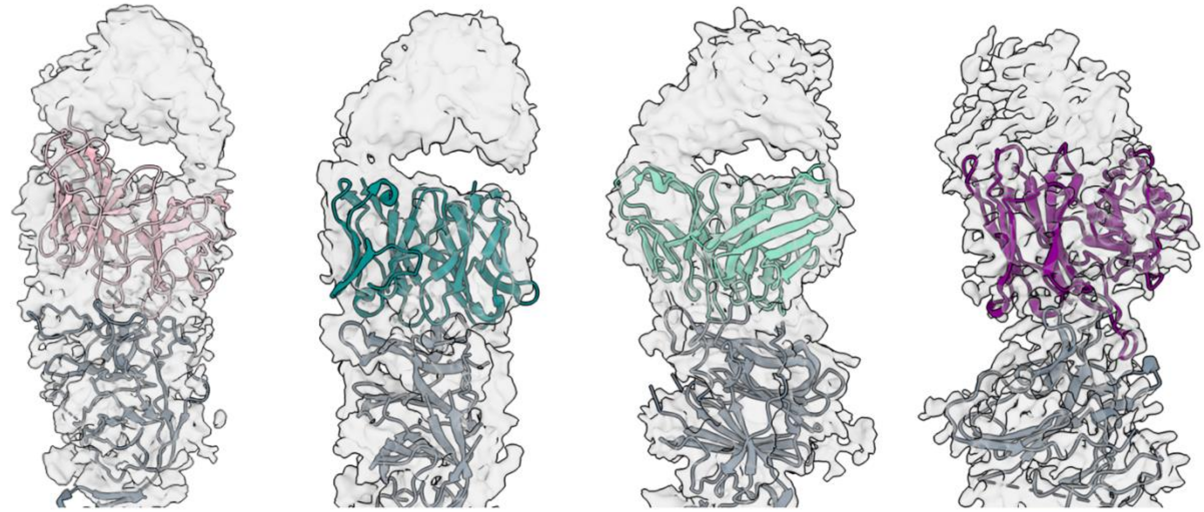


**Figure S9. Published monoclonal Spike-NTD antibodies resembling ModWu-NTD pAbs.** Atomic models of NTD in complex with mAbs 2-17 (PDB# 7LQW), mAb S2M28 (PDB# 7LY3), 4-18 (PDB# 7L2E), and TXG-0078 (PDB# 8SWH) docked into cryo-EMPEM map densities (transparent) of pAbs ModWu-NTD2, ModWu-NTD3, ModWu-NTD4, and ModWu-NTD5, respectively.

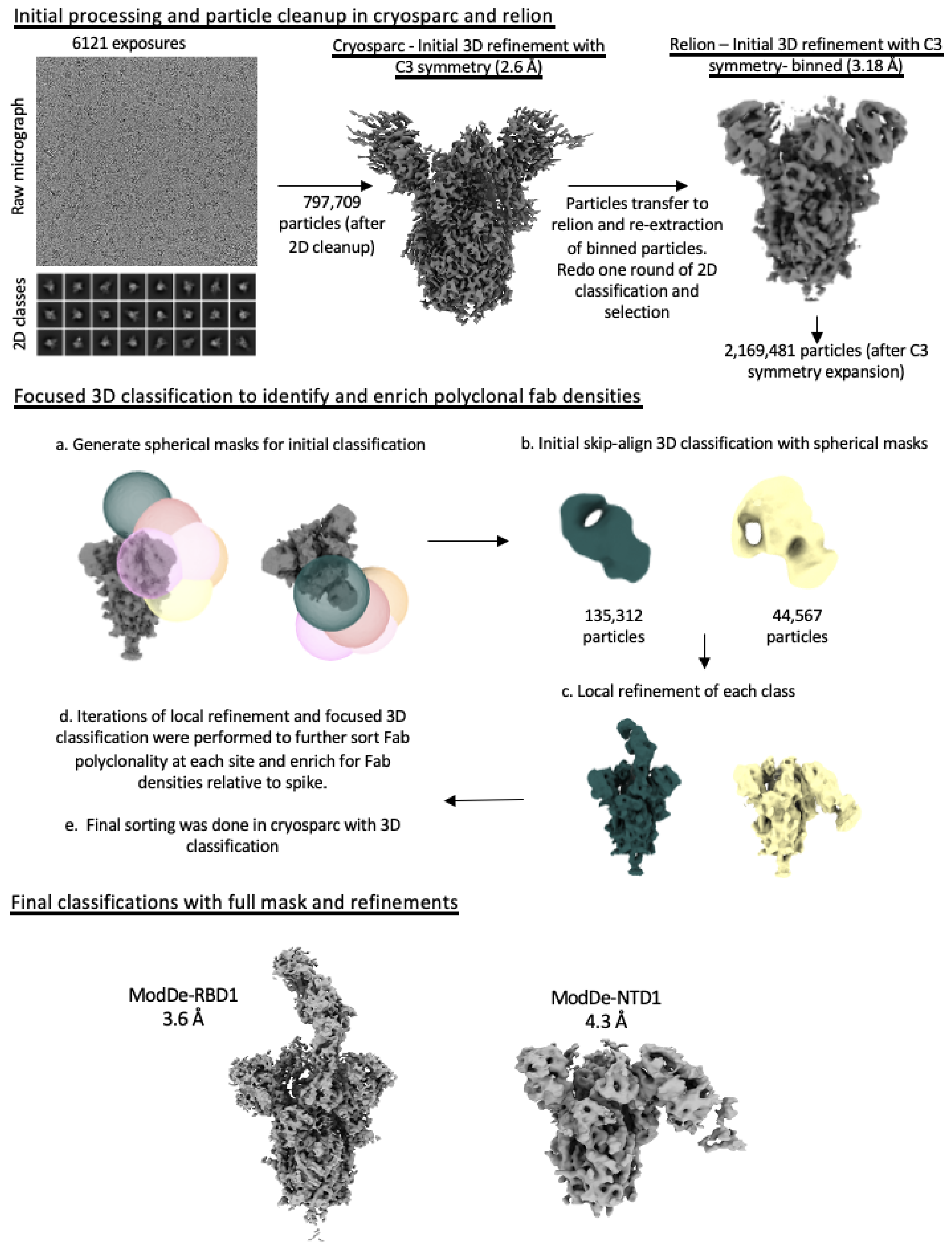

Initial processing and particle cleanup in cryosparc and relion
6121 exposures
Raw micrograph
2D classes
797,709 particles (after 2D cleanup)
Cryosparc - Initial 3D refinement with C3 symmetry (2.6 Å)
Particles transfer to relion and re-extraction of binned particles. Redo one round of 2D classification and selection
Relion – Initial 3D refinement with C3 symmetry- binned (3.18 Å)
2,169,481 particles (after C3 symmetry expansion)
Focused 3D classification to identify and enrich polyclonal fab densities
a. Generate spherical masks for initial classification
b. Initial skip-align 3D classification with spherical masks
135,312 particles
44,567 particles
c. Local refinement of each class
d. Iterations of local refinement and focused 3D classification were performed to further sort Fab polyclonality at each site and enrich for Fab densities relative to spike.
e. Final sorting was done in cryosparc with 3D classification
Final classifications with full mask and refinements
ModDe-RBD1
3.6 Å
ModDe-NTD1
4.3 Å

**Figure S10. Schematic representation of the cryo-EMPEM processing workflow for pooled mRNA-1273 vaccinated donor serum pAbs (day 43) complexed with SARS-CoV-2 Delta-HP-Mut7 Spike.**

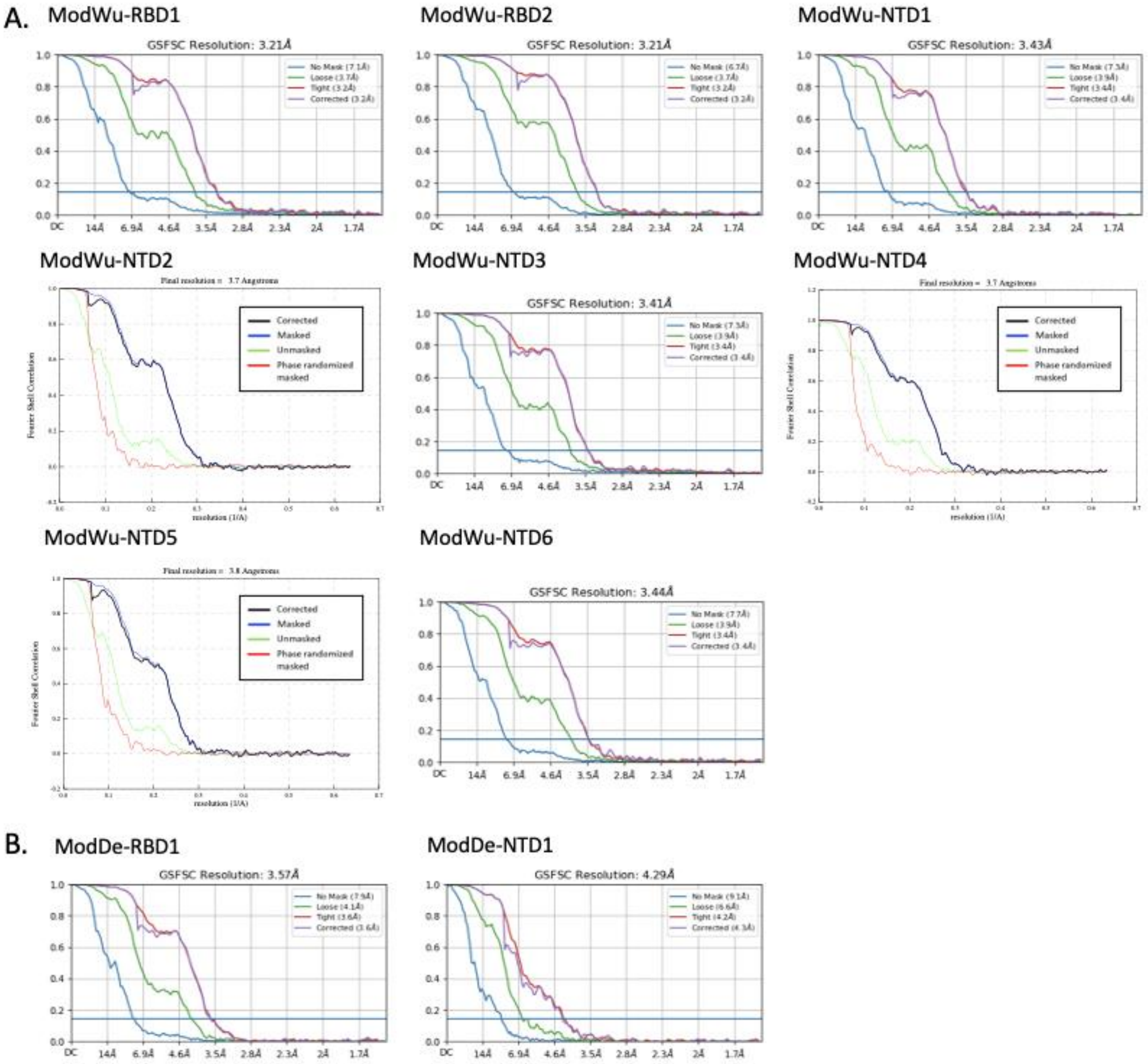


**Figure S11. FSC curves for mRNA-1273 vaccinated donor serum pAbs (day 43) complexed with (A) SARS-CoV-2-HP-GSAS-Mut7-D614G or (B) SARS-CoV-2 Delta-HP-Mut7 Spike**

#### Table S1. Cryo-EM data collection

| Data collection | NVX-NHPWu pAbs complexed with SARS-CoV-2-HP-GSAS-Mut7-D614G Spike | NVX-NHPSA pAbs complexed with SARS-CoV-2-HP-GSAS-Mut7-D614G Spike (2 sessions) | Mod-D49 pAbs complexed with SARS-CoV-2-HP-GSAS-Mut7-D614G Spike | Mod-D49 pAbs complexed with SARS-CoV-2 Delta-HP-Mut7 Spike |
|---|---|---|---|---|
| Microscope | FEI Titan Krios | FEI Titan Krios | FEI Titan Krios | FEI Titan Krios |
| Voltage (kV) | 300 | 300 | 300 | 300 |
| Detector | Gatan K2 Summit | Gatan K2 Summit | Gatan K3 | Gatan K3 |
| Recording mode | Counting | Counting | Super-resolution | Super-resolution |
| Nominal magnification | 130,000 | 130,000 | 29,000 | 29,000 |
| Movie micrograph pixel size (Å) | 1.045 | 1.045 | 0.788 | 0.788 |
| Number of frames per movie micrograph (Falcon 4 EER fractions) | 35 | 35 | 67 | 67 |
| Total dose ($e^-/Å^2$) | 50.6 | 50.6/50.1 | 60 | 60 |
| Number of movie micrographs | 6996 | 7942 | 14094 | 7958 |

#### Table S2. Cryo-EMPEM refinement parameters for NVX-NHPWu pAbs

| Map | NVX-NHPWu1 | NVX-NHPWu2 | NVX-NHPWu3 | NVX-NHPWu4 | NVX-NHPWu5 | NVX-NHPWu6 | NVX-NHPWu7 |
|---|---|---|---|---|---|---|---|
| EMDB | EMD-48199 | EMD-48200 | EMD-48201 | EMD-48202 | EMD-48203 | EMD-48204 | EMD-48205 |
| Number of molecular projection images in map | 13,248 | 13,989 | 12,190 | 12,140 | 11,441 | 13,125 | 37,096 |
| Symmetry | C1 | C1 | C1 | C1 | C1 | C1 | C1 |
| Map resolution (FSC 0.143; Å) | 4.2 | 4.2 | 4.2 | 4.3 | 4.3 | 4.4 | 3.8 |
| Map sharpening B-factor ($Å^2$) | -55.9 | -55.9 | -52.4 | -52 | -46.1 | -57.4 | -75.8 |

#### Table S3. Cryo-EMPEM refinement parameters for NVX-NHPSA pAbs

| **Map** | **NVX-NHPSA1** | **NVX-NHPSA2** | **NVX-NHPSA3** | **NVX-NHPSA4** | **NVX-NHPSA5** | **NVX-NHPSA6** | **NVX-NHPSA7** | **NVX-NHPSA8** |
|---|---|---|---|---|---|---|---|---|
| EMDB | EMD-48206 | EMD-48207 | EMD-48208 | EMD-48209 | EMD-48210 | EMD-48211 | EMD-48212 | EMD-48213 |
| Number of molecular projection images in map | 22,488 | 26,470 | 32,495 | 16,523 | 14,422 | 19,846 | 15,599 | 16,158 |
| Symmetry | C1 | C1 | C1 | C1 | C1 | C1 | C1 | C1 |
| Map resolution (FSC 0.143; Å) | 4.2 | 4.0 | 4.0 | 4.5 | 4.5 | 4.2 | 4.4 | 4.4 |
| Map sharpening B-factor ($Å^2$) | -72 | -67.6 | -70.3 | -85.1 | -85.9 | -70 | -85.5 | -88 |

#### Table S4. Cryo-EMPEM refinement parameters for ModWu pAbs

| **Map** | **ModWu-RBD1** | **ModWu-RBD2** | **ModWu-NTD1** | **ModWu-NTD2** | **ModWu-NTD3** | **ModWu-NTD4** | **ModWu-NTD5** | **ModWu-NTD6** |
|---|---|---|---|---|---|---|---|---|
| EMDB | EMD-48188 | EMD-48189 | EMD-48190 | EMD-48192 | EMD-48193 | EMD-48194 | EMD-48195 | EMD-48196 |
| Number of molecular projection images in map | 18,968 | 26,146 | 16,741 | 18,469 | 15,750 | 29,024 | 22,055 | 19,865 |
| Symmetry | C1 | C1 | C1 | C1 | C1 | C1 | C1 | C1 |
| Map resolution (FSC 0.143; Å) | 3.2 | 3.2 | 3.4 | 3.7 | 3.4 | 3.7 | 3.8 | 3.4 |
| Map sharpening B-factor ($Å^2$) | -46.8 | -56.1 | -66.2 | -6.25 | -42.4 | -3.1 | -84.8 | -44.4 |

### Table S5. Cryo-EMPEM refinement parameters for ModDe pAbs

| **Map** | **ModDe-RBD1** | **ModDe-NTD1** |
|---|---|---|
| EMDB | EMD-48197 | EMD-48198 |
| Number of molecular projection images in map | 15,351 | 8,325 |
| Symmetry | C1 | C1 |
| Map resolution (FSC 0.143; Å) | 3.6 | 4.3 |
| Map sharpening B-factor (Å$^2$) | -55.4 | -56 |
